# Multi-scale dynamics influence the division potential of stomatal lineage ground cells in *Arabidopsis*

**DOI:** 10.1101/2024.08.21.609020

**Authors:** Hannah F. Fung, Gabriel O. Amador, Renee Dale, Yan Gong, Macy Vollbrecht, Joel M. Erberich, Andrea Mair, Dominique C. Bergmann

## Abstract

During development, many precursor lineages are flexible, producing variable numbers and types of progeny cells. What factors determine whether a precursor cell differentiates or retains the capacity to divide? Here, we leverage the developmental flexibility of the stomatal lineage ground cell (SLGC) in *Arabidopsis* leaves as a model for how flexible decisions are regulated. Using a quantitative approach that combines long-term live imaging and statistical modeling, we discover that cell size is a strong predictor of SLGC behaviour: larger SLGCs divide less often than smaller cells. We propose that cell size is linked to division behaviour at multiple spatial scales. At the neighbourhood scale, cell size correlates with the strength of cell-cell signaling, which affects the rate at which SPEECHLESS (SPCH), a division-promoting transcription factor, is degraded. At the subcellular scale, cell size correlates with nuclear size, which modulates the concentration of SPCH in the nucleus. Our work shows how initial differences in SPCH levels are canalized by nuclear size and cell-cell signaling to inform the behaviour of a flexible cell type.

## Introduction

During development, different precursor lineages give rise to the full complement of cell types in a multicellular organism. Some lineages are more rigid, generating a fixed number of progeny of a certain type, while others are more flexible, producing variable numbers and types of progeny. The latter includes many plant lineages, which respond to environmental conditions to build tissues of different sizes and composition throughout an organism’s life. The *Arabidopsis* stomatal lineage offers a tractable system in which to investigate the emergence of stereotyped, but flexible patterns.

In the developing leaf, stomatal lineage cells undergo a series of asymmetric cell divisions (ACDs) that produce two daughter cells. The smaller daughter, called the meristemoid, can either differentiate into a guard mother cell (GMC) and ultimately a stoma, or divide asymmetrically one to five times before differentiating (Fig. 1A). The larger daughter, called the stomatal lineage ground cell (SLGC), faces a similar choice: it can either differentiate into a pavement cell or divide asymmetrically to generate a meristemoid and SLGC (Fig. 1A). SLGCs are often described as the larger “differentiating daughters”, destined to form pavement cells (e.g. Facette et al., 2019; Guo et al., 2021; Zhang et al., 2023), but this is a mischaracterization. SLGCs do divide asymmetrically, though at lower frequencies than meristemoids (Gong et al., 2021).

**Figure 1.**
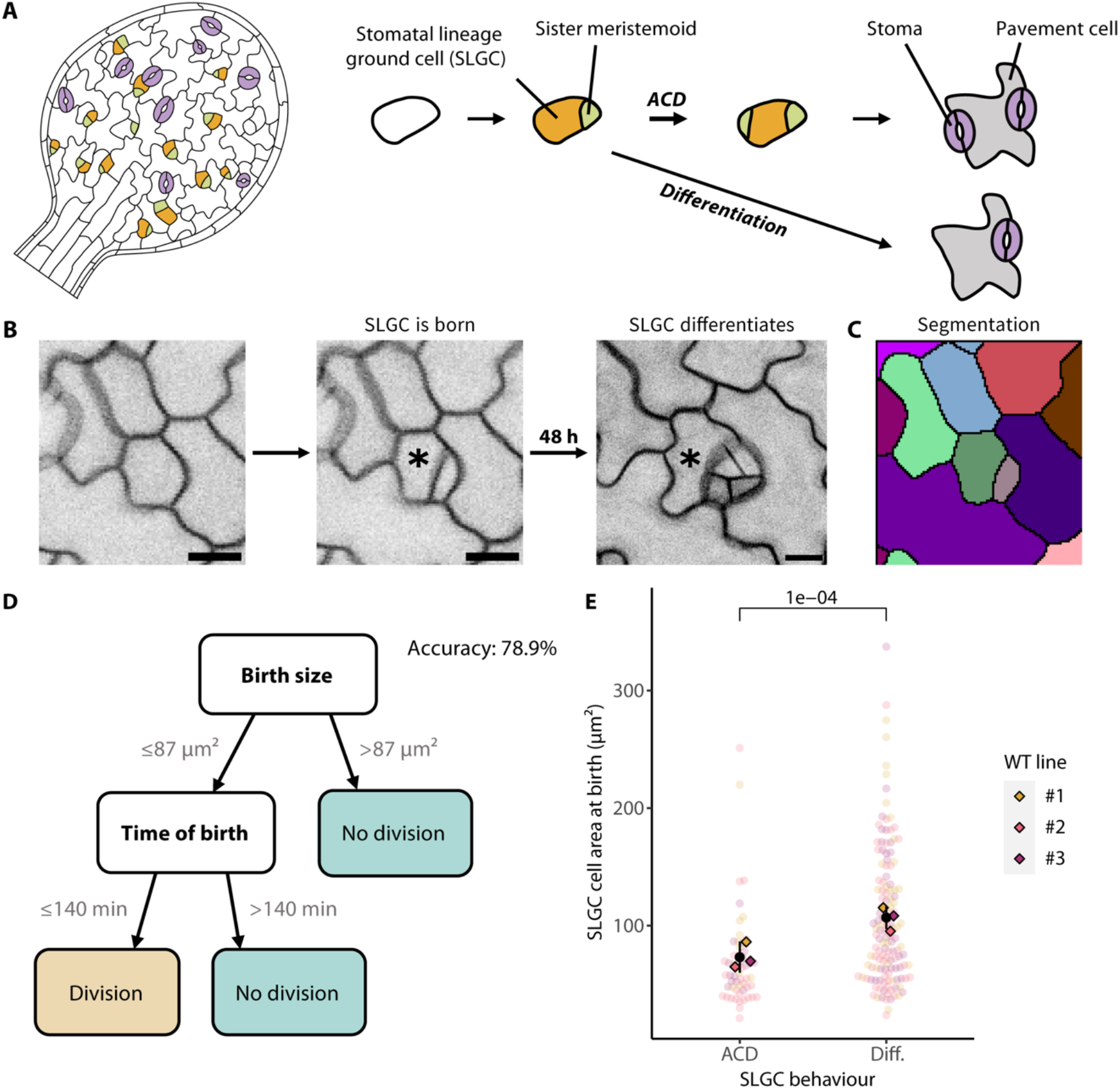
A decision tree identifies birth size as the strongest predictor of SLGC behaviour. (A) A cartoon of a developing *Arabidopsis* cotyledon (left), with a diagram of the stomatal lineage (right). Stomatal lineage cells divide asymmetrically to produce a smaller meristemoid (green) and a larger stomatal lineage ground cell (SLGC, gold). Meristemoids ultimately differentiate into stomata (purple). SLGCs can either divide asymmetrically (ACD) or differentiate into pavement cells (grey). (B) Illustration of the imaging-based approach. For each newly born SLGC (marked with an asterisk), we measured 15 features at birth. Two days later, we re-imaged the cell to capture its behaviour. In this example, the SLGC differentiated. Scale bar: 10 µm. (C) Cell segmentation of the second time point in (B), for semi-automated quantification of cellular features. (D) Decision tree following cost complexity pruning. Birth size was the strongest predictor of SLGC behaviour, followed by time of birth. (E) Cell areas at birth of SLGCs that divided (ACD) or differentiated (Diff.). Black circles and lines are individual-level means and standard deviations. The *P*-value is from a mixed-effects model with behaviour as a fixed effect and individual as a random effect. *N* = 3 individuals, 50-80 cells per individual. *Figure supplement:* Supplemental Figure 1.

A growing body of work suggests that SLGC divisions are actively suppressed. During asymmetric cell divisions, the mother cell segregates several polarity proteins to the SLGC, where they act as molecular scaffolds for a mitogen-activated protein kinase (MAPK) signaling cascade that promotes the degradation of SPEECHLESS (SPCH), a transcription factor required for asymmetric cell divisions (Dong et al., 2009; Guo et al., 2021; Houbaert et al., 2018; Rowe et al., 2019; Zhang et al., 2016, 2015). In contrast, SPCH levels remain high in the meristemoid sister, allowing it to divide multiple times before differentiating.

Although this model explains why meristemoids divide more often than SLGCs, it does not account for the flexibility of SLGC behaviours: why do some SLGCs divide, while others differentiate? This gap in knowledge is striking, given SLGCs have profound effects on the cell type composition of a leaf, responding to external inputs to generate three-quarters of all stomata (Geisler et al., 2000; Vatén et al., 2018). What are the factors that predispose an SLGC toward division or differentiation? Do the behaviours of individual SLGCs reflect their lineage history or their cellular neighbourhood?

Past efforts to characterize SLGCs have been hampered by the lack of cell-type-specific markers that distinguish SLGCs from their sister meristemoids in unequivocal ways. Consequently, it has been challenging to isolate SLGCs for transcriptomics (Ho et al., 2021) or to employ molecular techniques that rely on cell-type-specific promoters to manipulate cells. The subtle and quantitative phenotypes expected from the loss of regulators of SLGC behaviour make forward genetic screens infeasible.

Over the past two decades, quantitative studies have been instrumental in driving our understanding of processes that have eluded more traditional genetic approaches. For example, in the *Drosophila* embryo, careful *in vivo* measurements of the Bicoid transcription factor have offered fresh insight into the mechanisms by which morphogen gradients are established (reviewed by Saunders, 2021). Quantitative analyses have also overturned models: a recent study of the *Arabidopsis* root challenged a model of how formative cell divisions are regulated in the stem cell niche (Winter et al., 2024).

Here we developed a quantitative approach that combines long-term imaging and statistical modeling to identify correlates of SLGC behaviour at multiple spatiotemporal scales. Using this approach, we discovered that cell size is a strong predictor of SLGC behaviour: larger SLGCs divide less often than smaller cells. While we recently reported a size-based fate decision for other leaf epidermal cells (Gong, Dale, Fung, Amador et al., 2023), in this study we go further by providing a molecular explanation for the link between SLGC size and behaviour. We propose that cell size is linked to division behaviour at multiple spatial scales. At the neighbourhood scale, cell size correlates with the strength of cell-cell signaling, which affects the rate at which SPCH is degraded. At the subcellular scale, cell size correlates with nuclear size, which modulates the concentration of SPCH in the nucleus. Our work shows how initial differences in SPCH levels are canalized by nuclear size and cell-cell signaling to inform the behaviour of a flexible cell type.

## Results

### A decision tree identifies birth size as the strongest predictor of SLGC behaviour

Given the importance of SLGCs in leaf flexibility, but the challenges in using traditional genetic approaches to identify factors regulating this cell type, we adopted a holistic, imaging-based approach. We developed an imaging pipeline to measure cellular features at multiple scales, capturing SLGCs in an intact, wild-type, 3-day post germination (dpg) cotyledon from birth to their final, developmental outcomes (Fig. 1B). We then segmented the cell outlines using ilastik (Berg et al., 2019; Fig. 1C), which enabled the semi-automated quantification of 15 features at birth, including time of birth, tissue position, cell size and shape, mother identity, and characteristics of the immediate neighbours (Supplemental Fig. 1A). Two days later, we re-imaged the cotyledon and recorded whether each SLGC divided or differentiated (Fig. 1B). This two-day interval captures the vast majority of SLGC divisions (Supplemental Fig. 1B).

To pinpoint the most predictive features, we fed our measurements into a classification and regression tree (decision tree), which selects predictive features and orders them from most to least predictive. After cost complexity pruning to prevent overfitting (Supplemental Fig. 1C), we obtained a decision tree with a prediction accuracy of 78.9% (Fig. 1D). SLGC birth size was the strongest predictor, followed by time of birth (Fig. 1D). We also specified a random forest classifier which yielded very similar results: birth size had the highest feature importance score, followed by time of birth (Supplemental Fig. 1D).

To ensure these findings were robust, we captured and analyzed time-lapses of two other wildtype lines bearing different fluorescent reporters (see Methods). Across the three lines, size was consistently predictive of behaviour: only smaller cells could divide again, while larger cells differentiated (Fig. 1E). Surprisingly, this is the opposite of what we see in meristemoids, where the probability of dividing asymmetrically increases with cell size (Gong, Dale, Fung, Amador et al., 2023).

### Larger cells are born with lower SPCH concentrations

To understand why the relationship between size and behaviour is inverted in SLGCs (relative to meristemoids), it is useful to identify the specific genes or proteins involved. We therefore turned our attention to one of the few well-characterized proteins present in SLGCs, the transcription factor SPCH. Previous work reported that the frequency of SLGC divisions increased upon cytokinin signalling manipulations, and that the SLGCs expressed SPCH before dividing (Vatén et al., 2018). Whether SPCH has a similar role during normal development, however, was not determined. Nevertheless, the cytokinin results provide testable hypotheses about the relationship between SPCH levels and size-dependent divisions, namely that (1) SPCH levels should correlate with SLGC behaviour; and (2) larger cells should contain less SPCH.

To quantify SPCH levels, we captured timelapses of 3-dpg cotyledons expressing a translational reporter (*pSPCH::SPCH-YFP* rescuing *spch-3-/-*; Lopez-Anido et al., 2021; Gong, Dale, Fung, Amador et al., 2023) and tracked SPCH intensities from cell birth to the end of the time-lapse (Fig. 2A-B). SPCH was exclusively nuclear during this period. Surprisingly, SPCH intensities were already predictive at birth: dividing cells were born with significantly more SPCH than differentiating cells (Fig. 2C).

**Figure 2.**
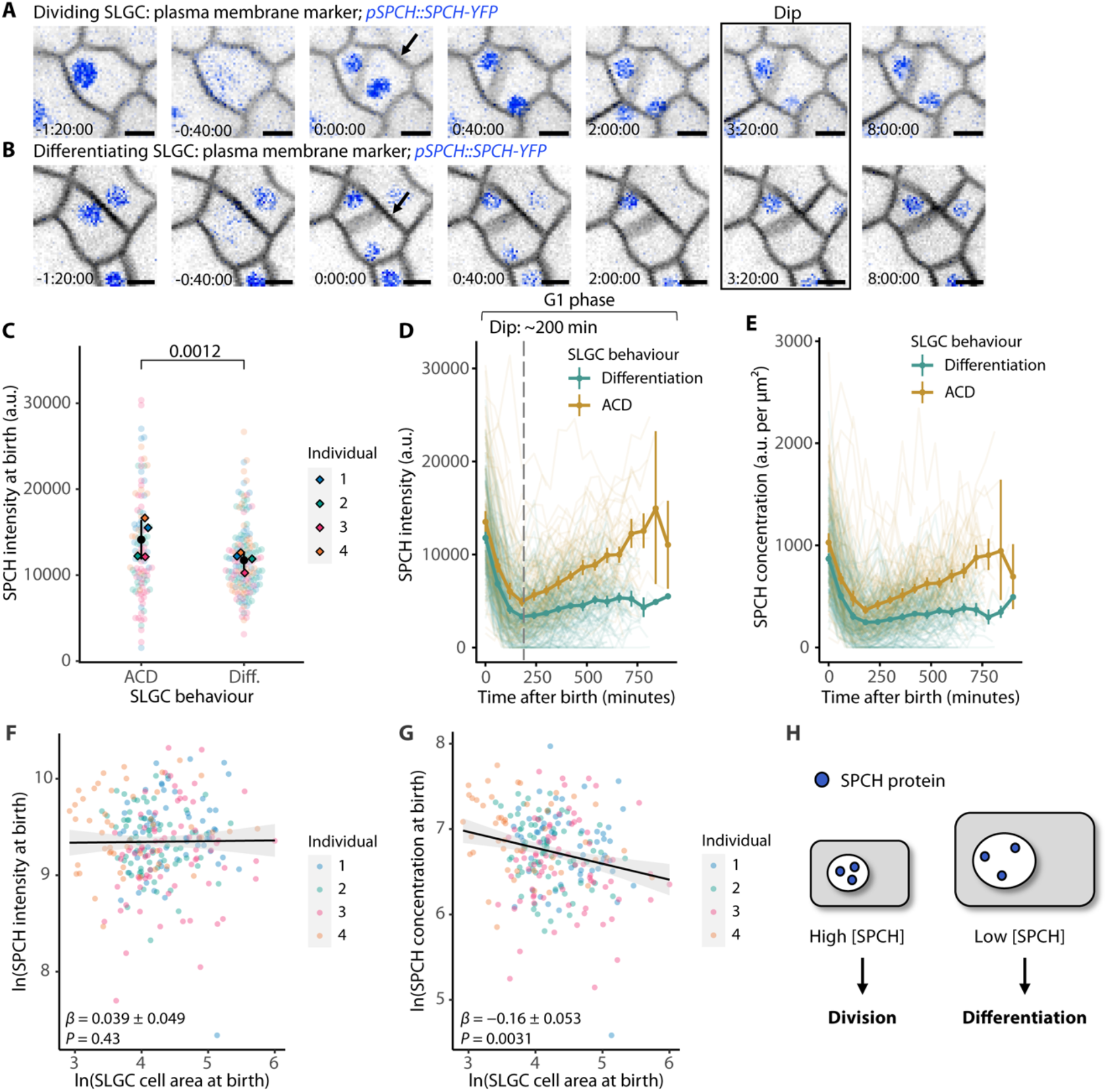
Larger cells are born with lower concentrations of SPCH and divide less often. (A-B) Inverted confocal images of the SPCH translational reporter *pSPCH::SPCH-YFP; spch-3* in a dividing cell (A) or a differentiating cell (B) at 3-dpg. The SLGCs were born at 0 minutes (arrows). Scale bar: 5 µm. (C) SPCH intensities at birth in dividing (ACD) or differentiating (Diff.) cells. Black circles and lines are individual-level means and standard deviations. (D-E) SPCH intensities over time (D) or SPCH nuclear concentrations over time (E), coloured by behaviour. Circles and vertical lines are binned means and bootstrapped 95% confidence intervals. (F-G) SPCH intensity at birth (F) or SPCH nuclear concentration at birth (G) vs. cell area at birth. Axes are ln-transformed. Black lines and grey bands are linear model predictions and 95% confidence intervals. (H) A cartoon of the SPCH dilution model. Small and large cells are born with comparable SPCH intensities, a proxy for the number of SPCH molecules. Because larger cells have larger nuclei, they are born with lower concentrations of SPCH and divide less often. (C,F-G) *P*-values are from mixed-effects models with individual as a random effect. (C-G) *N* = 4 individuals, 50-75 cells per individual. *Figure supplements:* Supplemental Figure 2, Supplemental Table 1.

After birth, SPCH intensities declined dramatically for ∼200 minutes in both dividing and differentiating populations (Fig. 2D). While repeated imaging can lead to photobleaching and a decline in SPCH intensities, we confirmed that bleaching alone could not account for a decline of this magnitude (see Methods; Supplemental Table 1). At ∼200 minutes after birth, SPCH levels began to rise in dividing, but not in differentiating cells (Fig. 2D).

To determine whether the rise in SPCH levels was a consequence of cell cycle progression, we compared SPCH dynamics to that of the replication licensing factor CDT1A, which accumulates during G1 and is rapidly degraded at the G1/S transition (Desvoyes et al., 2019). The rise in SPCH intensities preceded that of CDT1A by ∼100 minutes (Supplemental Fig. 2A), implying that SPCH is a cause, rather than a consequence of cell cycle progression. The dynamics of SPCH nuclear concentration over time (SPCH intensity divided by nuclear area) resembled those of SPCH intensity (Fig. 2E).

Next, we tested whether larger SLGCs contain less SPCH. There are two ways of measuring the ‘amount’ of SPCH in a cell: fluorescence intensity, which scales with the number of molecules of SPCH, and nuclear concentration, which also accounts for nuclear size. Of the two measures, concentration is likely more biologically meaningful because it contributes directly to transcription by affecting binding site occupancy (Doughty et al., 2024). Although we did not detect a significant correlation between SPCH intensity and cell size at birth (Fig. 2F), we found that larger cells were born with lower nuclear concentrations of SPCH (Fig. 2G). Since larger cells have larger nuclei (Supplemental Fig. 2B), these data suggest that SPCH is diluted in larger cells (Fig. 2H). Taken together, our results indicate that larger cells are born with lower concentrations of SPCH, which may explain why they divide less often.

### Large SLGCs divide more often when SPCH levels are increased

A SPCH “dilution” hypothesis also predicts that large SLGCs will divide more often if their SPCH concentration is increased. To address this prediction, we first determined whether large cells are capable of responding to a general division-promoting factor. We expressed the D-type cyclin CYCLIN D7;1 (CYCD7;1) under the epidermis-specific *ATML1* promoter (*pATML1::CYCD7;1-YFP*; Weimer et al., 2018) in wild-type cotyledons. The construct was expressed in all epidermal cells at 3-dpg (Supplemental Fig. 3A) and induced both small and large SLGCs to divide (Supplemental Fig. 3B-C). Thus, we can conclude that large SLGCs are division-competent.

Next, we tested whether large cells divide more often when supplied with more SPCH. We generated a *pSPCH::SPCH-YFP; spch-3* line where SPCH was overproduced in its normal expression domain (SPCH++; see Methods). Across cell sizes, SPCH intensities were higher in SPCH++ cotyledons than in those expressing the SPCH translational reporter (Supplemental Fig. 3D). Accordingly, we observed an increase in the proportion of SLGCs that divided, relative to wild-type cotyledons (Fig. 3A). This increase in division frequency was due to elevated SPCH levels, rather than a significant change in birth size (Fig. 3B).

**Figure 3.**
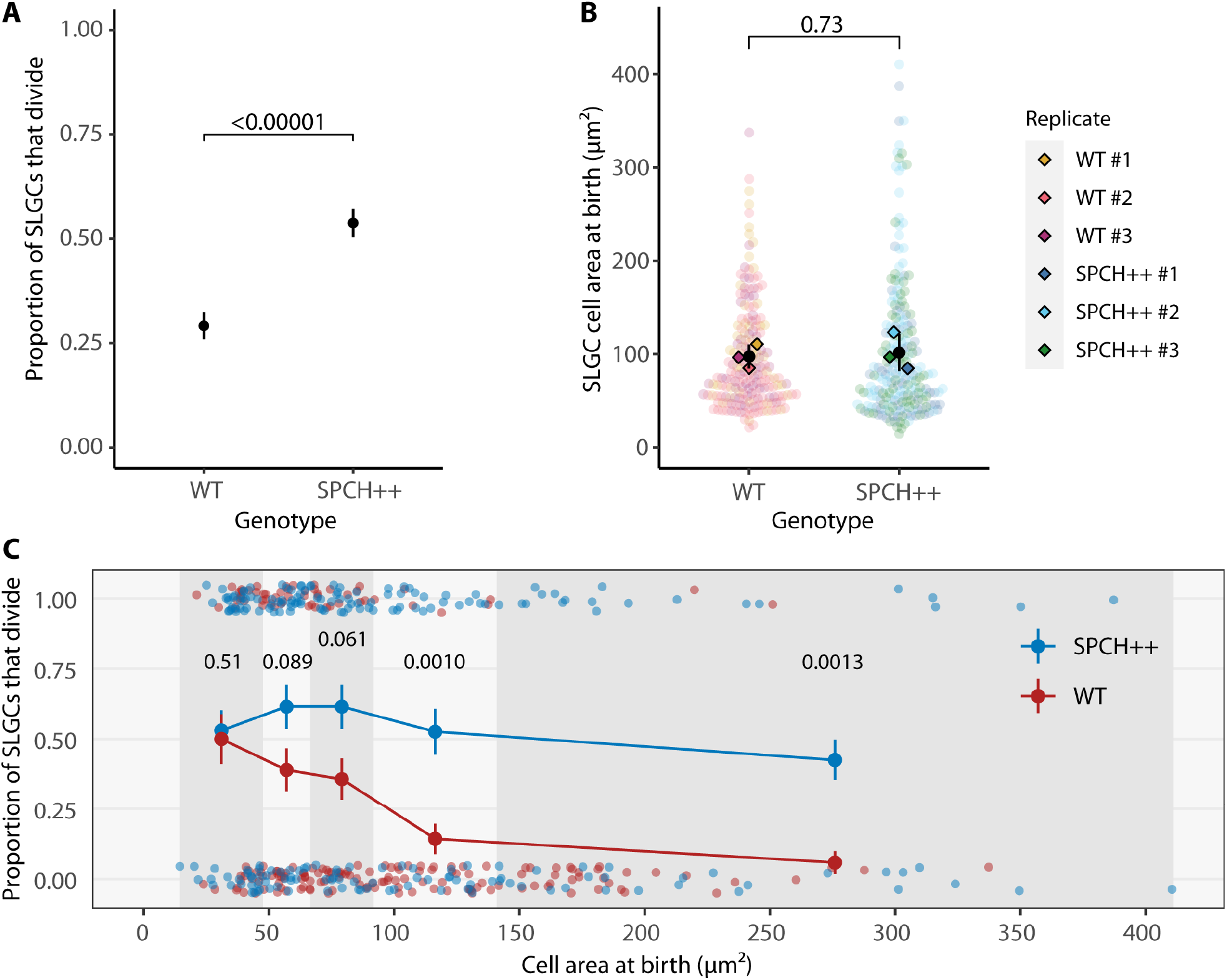
Large SLGCs divide more often when SPCH levels are increased. (A-B) The proportion of SLGCs that divided (A) or SLGC cell areas at birth (B) in wild-type seedlings and seedlings where SPCH accumulates to higher levels (strong *pSPCH::SPCH-YFP spch-3*; SPCH++). Black circles and lines are means and standard deviations. *P*-values are from a *Z*-test for two proportions (A) or a mixed-effects model with individual as a random effect (B). (C) The proportion of SLGCs that divided in wild-type and SPCH++ cotyledons. Vertical shading delineates quintiles of SLGC cell area at birth, from left: smallest 20%, 20-40%, 40-60%, 60-80%, largest 20%. Circles and lines are means and standard deviations. *P*-values are from Holm-Bonferroni corrected *Z*-tests. (A-C) *N* = 3 individuals per genotype, 50-80 cells per individual. *Figure supplement:* Supplemental Figure 3.

To determine specifically whether large cells divide more often when SPCH is overproduced, we binned wild-type and SPCH++ cells into quintiles based on size. For each quintile, we computed the proportion of cells that divided per genotype (Fig. 3C). Consistent with our hypothesis, SPCH++ cells divided more often than wild-type cells in the fourth and fifth quintiles, indicating that *SPCH* overexpression is sufficient to increase the proportion of large cells that divide (Fig. 3C).

### Number of signaling neighbours can influence SPCH degradation rates

One inescapable observation in the time-lapse imaging of SPCH is that this factor is highly dynamic. SPCH intensities decline after birth in all SLGCs (Fig. 2D), but interestingly, they appear to fall faster in differentiating cells. This prompted us to test whether the rate of SPCH decline is correlated with SLGC behaviour. We fit an exponential decay function (*N*(t) = *N*_0_ * *e*^λt^) to the SPCH intensities of each cell from 0 to ∼200 minutes after birth and estimated the decay constant λ (the more negative the constant, the higher the rate of decline). The decay constant was predictive of SLGC behaviour: SPCH levels fell more rapidly in differentiating cells than in dividing cells (Supplemental Fig. 4A). Among cells where SPCH levels declined, larger cells showed higher rates of decline than smaller cells (Supplemental Fig. 4B).

Why would SPCH levels decline faster in larger cells? Previous studies have shown that SPCH is regulated by the peptides EPIDERMAL PATTERNING FACTOR 1 and 2 (EPF1/2; Lee et al., 2015, 2012), which activate a MAPK signaling cascade that targets the SPCH protein in neighbouring cells for degradation (Lampard, MacAlister et al., 2008; Fig. 4A). EPF1 and EPF2 are reported to be secreted by meristemoids, GMCs, and young stomata to prevent their neighbours from developing into stomata (Hara et al., 2009, 2007; Hunt and Gray, 2009; Lee et al., 2012). This ensures that stomata are spaced apart, which optimizes stomatal function (Dow et al., 2014; Harrison et al., 2020). Although mobile peptides could act over large spatial scales, lineage tracing in stomatal signaling mutants (Geisler et al., 2000) suggests that the signals that establish and maintain stomatal spacing are likely juxtacrine.

**Figure 4.**
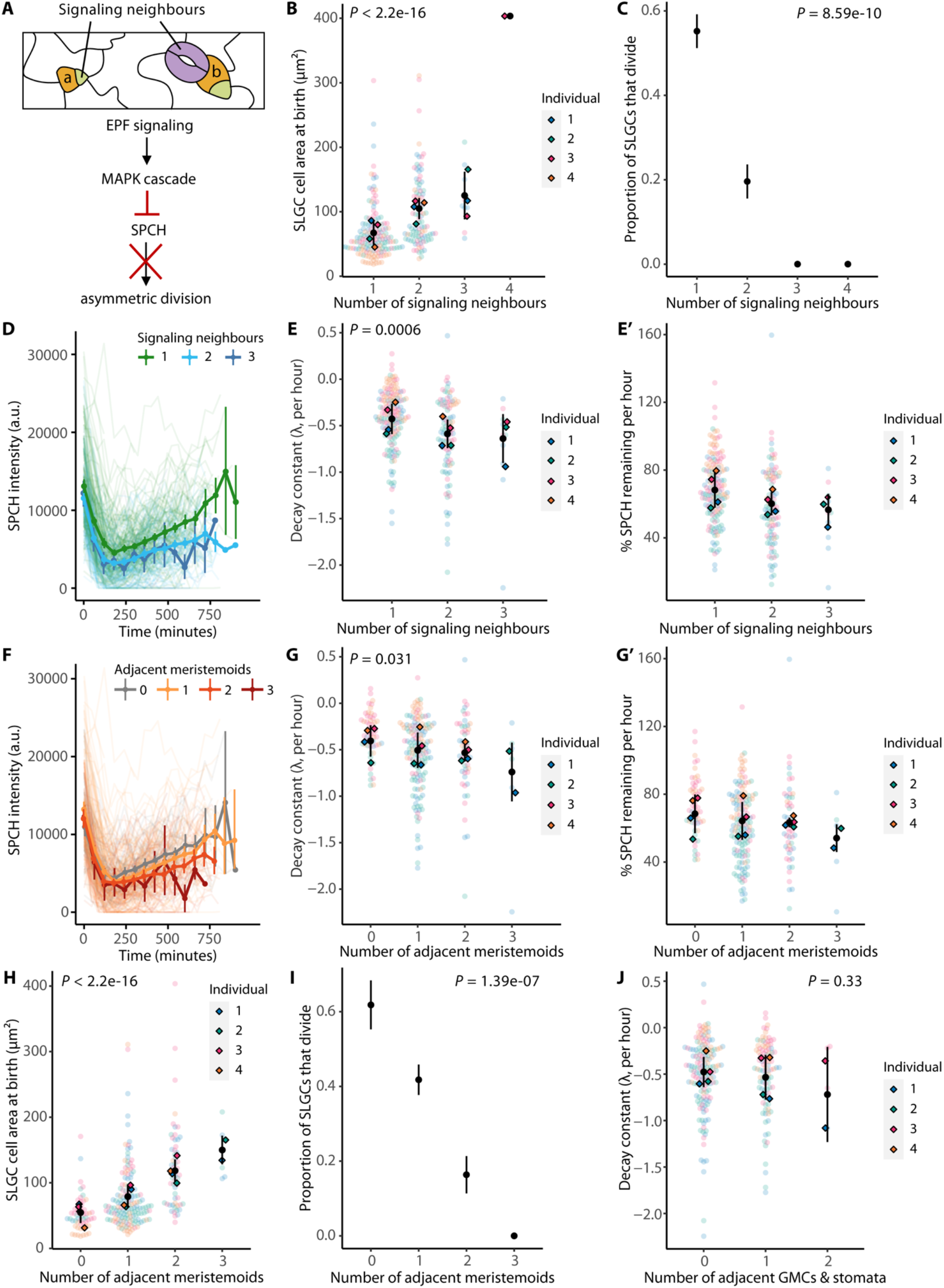
Cells with more signaling neighbours experience higher SPCH degradation rates and divide less often. (A) Cartoon of cells monitored in this figure (top), and diagram of the EPIDERMAL PATTERNING FACTOR (EPF) signaling cascade that targets SPCH for degradation (bottom). a: SLGC with one signaling neighbour. b: SLGC with two signaling neighbours. (B-C) SLGC cell area at birth (B) or the proportion of SLGCs that divided (C) by the number of signaling neighbours. (D) SPCH intensities over time, coloured by the number of signaling neighbours. (E) The decay constant, a measure of how fast SPCH is degraded, by the number of signaling neighbours. The more negative the constant, the higher the degradation rate. (E′) The same data, re-plotted as the percentage of SPCH remaining after every hour. SPCH levels declined in most cells, but we did observe some cells where SPCH increased (points >100%). (F) SPCH intensities over time, coloured by the number of adjacent meristemoids. (G-I) The decay constant (G), the percentage of SPCH remaining after every hour (G′), cell area at birth (H), or the proportion of SLGCs that divided (I) by the number of adjacent meristemoids. (J) The decay constant by the number of adjacent GMCs and stomata. (B,C,E,G-J) Black circles and lines are means and standard deviations. *P*-values are from mixed-effects models with individual as a random effect (B,E,G,H,J) or chi-squared tests for trend in proportions (C,I). (D,F) Circles and vertical lines are binned means and bootstrapped 95% confidence intervals. (B-J) *N* = 4 individuals, 50-75 cells per individual. *Figure supplement:* Supplemental Figure 4.

In light of the known EPF-MAPK signaling pathway, a plausible explanation for why larger SLGCs experience higher SPCH degradation rates is that they have more “signaling neighbours” (neighbours that are meristemoids, GMCs, or stomata; Fig. 4A). This geometric argument predicts that (1) cells with more signaling neighbours should be larger; (2) they should divide less often; and (3) they should experience higher SPCH degradation rates.

To test these predictions, we analyzed the time-lapses of 3-dpg cotyledons expressing the SPCH translational reporter used in Fig. 2. Cells with more signaling neighbours were larger (Fig. 4B) and divided less often (Fig. 4C). The number of signaling neighbours appeared to be a good proxy for the strength of EPF signaling, as neither the total number of neighbours nor the fraction of the cell perimeter in contact with a signaling neighbour was predictive of SLGC behaviour, after accounting for the number of signaling neighbours (Supplemental Fig. 4C-D). In line with our third prediction, cells with more signaling neighbours experienced higher SPCH degradation rates (Fig. 4D-E′).

These data conform to textbook “lateral inhibition” models in which mature stomata generate inhibitory fields to prevent the formation of adjacent stomata (reviewed in Nadeau and Sack, 2002). However, a closer look at our data separated by cell type reveals that meristemoids are largely responsible for the neighbour effect. Cells with more meristemoid neighbours experienced higher SPCH degradation rates (Fig. 4F-G′). They were also larger (Fig. 4H) and divided less often (Fig. 4I).

In contrast, neither the number of adjacent stomata nor the number of adjacent GMCs was significantly associated with SPCH degradation rates (Fig. 4J; Supplemental Fig. 4E-F). Very few of the SLGCs at 3-dpg had a stomatal neighbour, compromising our ability to estimate the mean SPCH degradation rate in this group of cells. Consequently, our data lacked the statistical power to detect an association (if any) between the number of stomata and degradation rate. Our GMC results, on the other hand, were not limited by statistical power. Here we considered two explanations for the lack of GMC influence; first, that GMCs do not suppress SLGC divisions, or second, that GMCs suppress SLGC divisions independently of SPCH degradation. Contrary to expectations from lateral inhibition models, cells with more GMC neighbours tended to divide more often (Supplemental Fig. 4H).

Finally, we tested whether the link between size and behaviour is abrogated when the ability of SPCH to respond to MAPK signaling is disrupted. In seedlings where *spch-3* is rescued with a SPCH variant lacking three MAPK phosphorylation sites (*pSPCH::SPCH2-4A-YFP;* Davies and Bergmann, 2014), many large SLGCs (>150 µm^2^) divided (Fig. 5A). In fact, dividing cells were slightly but significantly larger than differentiating cells (Fig. 5B), consistent with MAPK signaling suppressing the division potential of large SLGCs.

**Figure 5.**
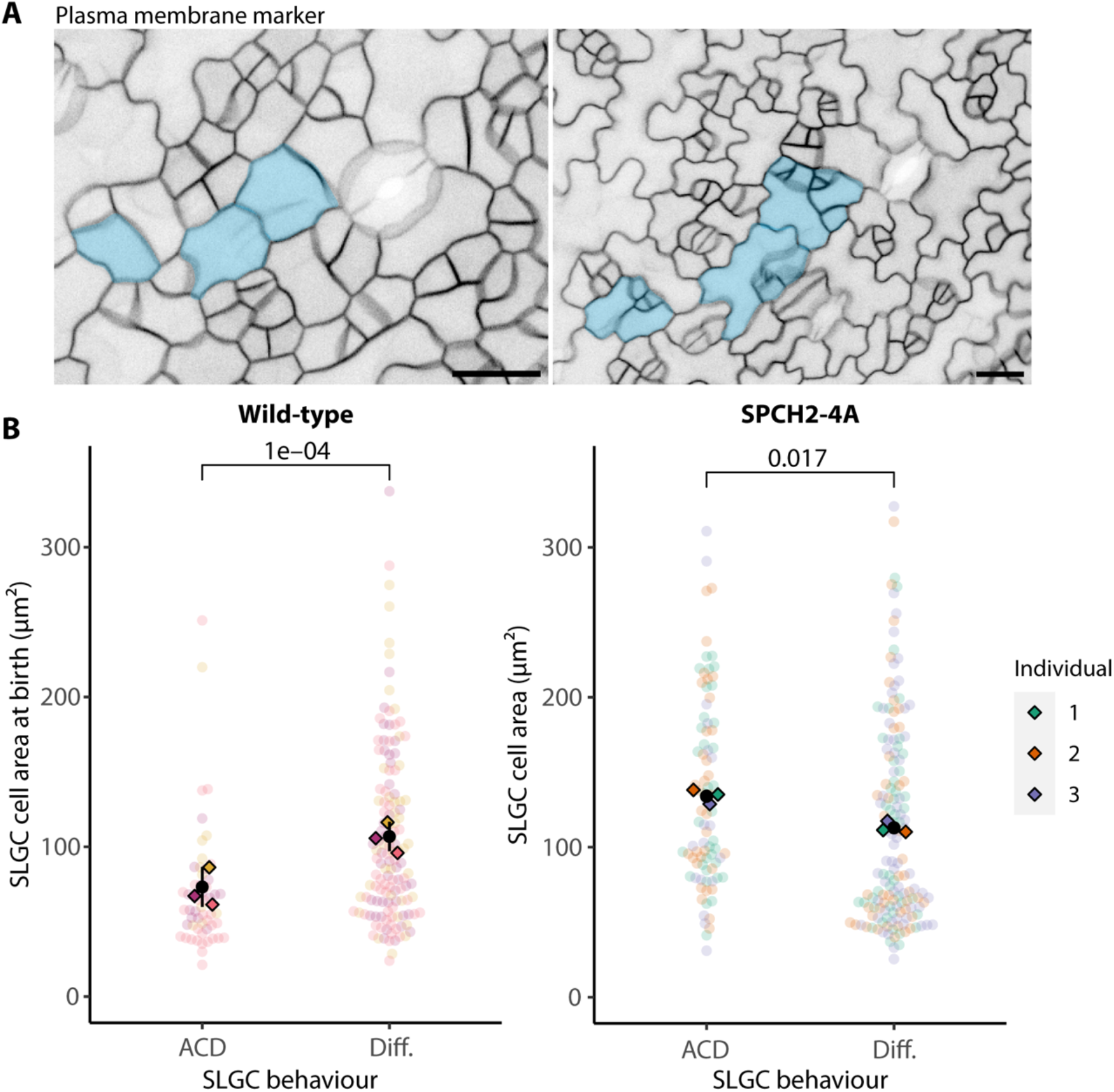
The link between size and behaviour is abrogated when the ability of SPCH to respond to MAPK signaling is disrupted. (A) Micrograph of a 3-dpg *spch-3* cotyledon expressing a SPCH variant lacking three MAPK phosphorylation sites (*pSPCH::SPCH2-4A-YFP*; left). The same region two days later (right). Large, dividing SLGCs are false-coloured in blue. Scale bar: 20 µm. (B) Cell areas of SLGCs that divided (ACD) or differentiated (Diff.) in wild-type and *pSPCH::SPCH2-4A-YFP; spch-3* (SPCH2-4A) cotyledons. The wild-type data are re-plotted from Fig. 1E. *N* = 3 individuals per genotype, 50-80 cells per individual.

### Simulations and statistical evaluations of models

So far, we have proposed that large cells divide less often because they have more signaling neighbours (pathway #1; Fig. 6A) and the SPCH they contain is diluted because of their larger nuclei (pathway #2; Fig. 6A). In these cells, the concentration of SPCH ([SPCH]) remaining after degradation (at the ‘dip’) is too low to activate the proposed positive feedback loop required to drive cell cycle progression (Horst et al., 2015; Lau et al., 2014). Our model yields two predictions that can be tested statistically: [SPCH] at the dip should be predictive of SLGC behaviour; and among cells with the same SPCH intensities at the dip, cells with larger nuclei should divide less often.

**Figure 6.**
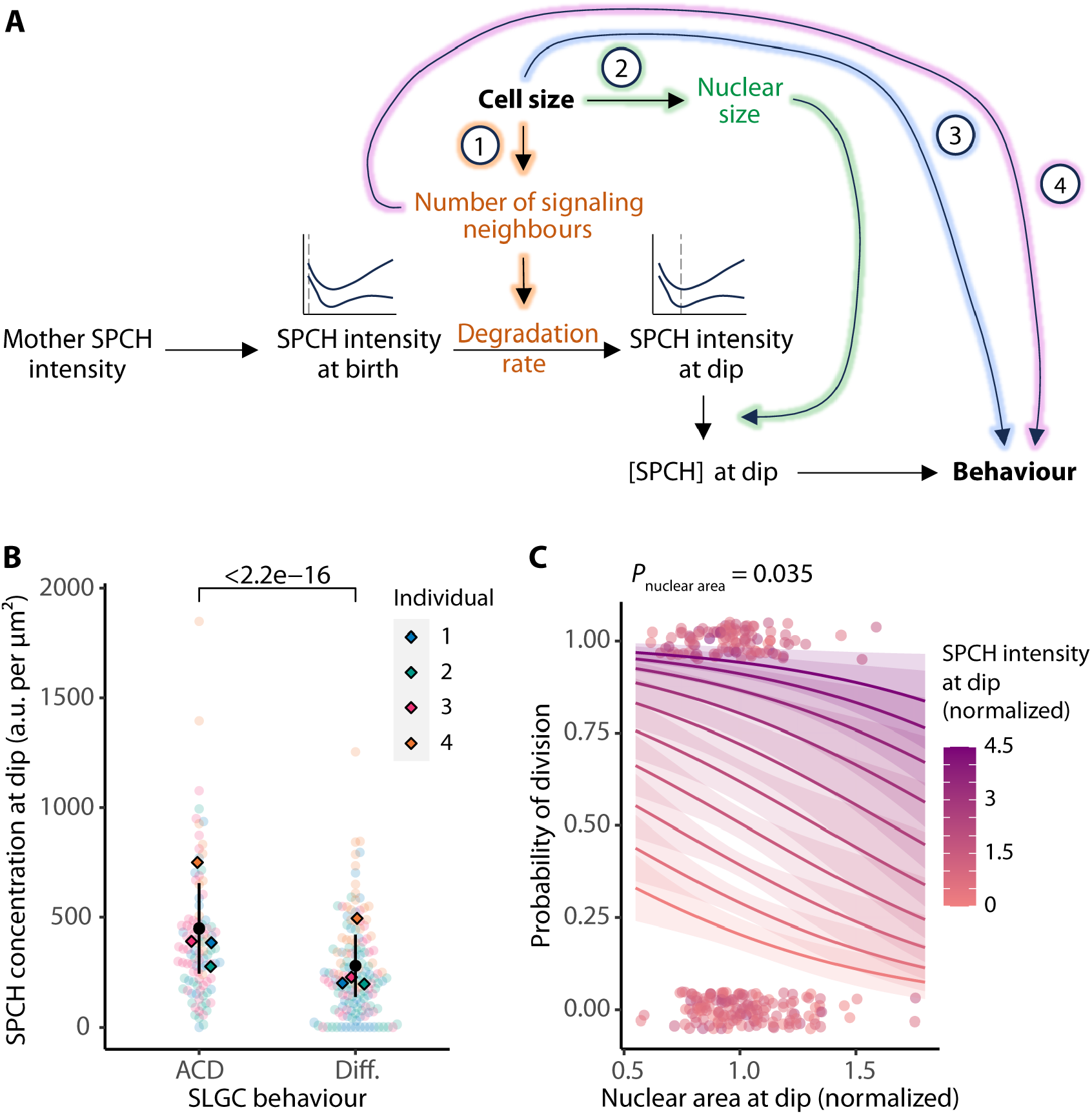
Size is linked to SLGC behaviour through SPCH degradation, SPCH concentration, and a SPCH-independent pathway. (A) Model for SLGC division propensity. Cell size is linked to SLGC behaviour through (1) SPCH degradation, gold; (2) SPCH concentration, green; and (3) a SPCH-independent pathway, blue. (4) Based on our simulations (Supplemental Figs 5-7; Supplemental Tables 2-4), signaling neighbours may affect behaviour in a SPCH- and size-independent manner, purple. (B) SPCH nuclear concentrations at the dip in dividing (ACD) or differentiating (Diff.) cells. The *P*-value is from a mixed-effects model with individual as a random effect. (C) Multiple logistic regression of the probability of division on nuclear area at the dip, controlling for SPCH intensity at the dip. Among cells with the same SPCH intensity at the dip, cells with larger nuclei divided less often, because the SPCH they contained was diluted by larger nuclear compartments. Data from different individuals were normalized (relative to the individual mean) and pooled for visualization. Lines and bands are logistic model predictions and standard errors. The *P*-value is from a logistic regression with individual as a blocking variable. (B-C) *N* = 4 individuals, 50-75 cells per individual. *Figure supplement:* Supplemental Figures 5-7; Supplemental Tables 2-4.

In line with the first prediction, dividing cells had significantly higher [SPCH] at the dip than differentiating cells (Fig. 6B). Consistent with the second prediction, cells with larger nuclei divided less often than those with smaller nuclei, after controlling for SPCH intensity at the dip (Fig. 6C). Both pathways in our model (Fig. 6A) converge on [SPCH] at the dip, which we hypothesize is a primary determinant of SLGC behaviour.

Is [SPCH] at the dip sufficient to recapitulate SLGC behaviours? To test this, we specified a stochastic and asynchronous rule-based lineage decision tree model, with [SPCH] at the dip as the sole determinant of behaviour (Supplemental Fig. 5; model details in methods). The model begins with a population of 1,000 mother cells, each with a randomly drawn size, SPCH intensity, and number of signaling neighbours. The mother cells divide with a randomly drawn asymmetry, producing a smaller meristemoid and a larger SLGC. Based on the measurements in Supplemental Fig. 6, each SLGC is assumed to inherit two-thirds of its mother’s SPCH intensity and to have one signaling neighbour more than its mother (i.e. the newly generated sister meristemoid).

After birth, SPCH is degraded according to one of four modes: the decay constants (λ) are randomly drawn; modulated by size; modulated by signaling neighbours; or modulated by both size and signaling neighbours. The SLGCs then undergo a fate-determining program in which cells with higher [SPCH] have a higher chance of dividing. We derived all input parameters by fitting theoretical distributions or logistic regressions to empirical measurements from four individual plants (see Supplemental Tables 2 and 3 for fitted parameters).

We assessed each model according to its ability to recapitulate the sizes and [SPCH] of dividing and differentiating cells (Supplemental Fig. 7A-B), and the proportion of cells that divided given the number of signaling neighbours (Supplemental Fig. 7C, Supplemental Table 4; see Methods for details). The highest-ranking model was one where degradation rates scaled with both size and signaling neighbours. However, it struggled to reproduce the negative relationship between signaling neighbours and behaviour (Supplemental Fig. 7C), suggesting that [SPCH] alone is insufficient to recapitulate the observed patterns.

To determine whether other features were required for the fate-determining process, we specified fate-determining programs that considered different combinations of cell size, [SPCH], signaling neighbours, and/or their interaction terms. The top model was one where degradation rates scaled with neighbours, and where size, [SPCH], and neighbours were all considered in the fate-determining process (Supplemental Fig. 7). The fact that a fate-determining program with size, [SPCH], and neighbours outperformed one with [SPCH] alone implies that there are SPCH-independent pathways through which size and neighbours operate (Fig. 6A).

## Discussion

In this study, we leverage quantitative approaches to define the properties and behaviours of the enigmatic stem-cell-like SLGCs in the *Arabidopsis* leaf epidermis. We show that the division potential of an SLGC is a product of its neighbourhood and nuclear size. Larger cells divide less often because they have more signaling neighbours and larger nuclei, resulting in lower [SPCH]. Through computational simulations, we also identify SPCH-independent pathways that may link cell size and signaling to behaviour.

In a previous study, we showed that cell size regulates meristemoid behaviour: larger meristemoids divide more often than smaller ones (Gong, Dale, Fung, Amador et al., 2023). Our findings here extend that work in two important ways. First, we make the surprising observation that while cell size is also associated with SLGC behaviour, the direction of this relationship is inverted: larger SLGCs divide *less* often than smaller ones. Second, by linking cell size to the dynamics of SPCH, we can propose a molecular explanation for why large SLGCs divide less often. More broadly, our work expands on existing studies of cell size and behaviour (e.g. Gong, Dale, Fung, Amador et al., 2023; Hubatsch et al., 2019; Kirk et al., 1993), which have focused primarily on cell-autonomous factors, to consider the neighbourhood of a cell. We demonstrate that cell size can affect non-cell-autonomous factors, such as the strength of cell-cell signaling.

There is a growing recognition of the importance of cell geometries for signaling (Fiorentino and Scialdone, 2022; Haftbaradaran Esfahani and Knöll, 2020). For example, Pentinmikko et al. (2022) used *in vitro* organoids and culture scaffolds to show that the area of neighbour contacts in small intestinal stem cells affects the strength of signals they receive. In these stem cells, apical constriction increases the lateral surface-to-volume ratio, which enhances their ability to receive niche signals from neighbouring Paneth cells. When the lateral surface-to-volume ratio was reduced, the stem cells initiated fewer organoids, suggesting their regenerative capacity was disrupted (Pentinmikko et al., 2022). Similarly, we report here that the size of an SLGC can affect the magnitude of the signals it receives, as measured by the rate of SPCH degradation.

In the leaf epidermis, stomata and their precursors (meristemoids and GMCs) are thought to secrete mobile peptides to prevent their neighbours from dividing asymmetrically and producing stomata. This ensures that stomata are spaced apart, which optimizes stomatal opening and environmental responsiveness (Dow et al., 2014). Unlike meristemoids, stomata and GMCs are committed cells that cannot respond to an adjacent stomatal precursor by dividing asymmetrically. Thus, one would expect the strongest inhibitory signals to come from these cells. Surprisingly, we find that meristemoids have the strongest effect on SLGC division potential among the three cell types. Signaling among progenitors may be critical to regulate cell numbers during the proliferative phase of leaf development. It is also possible that we have underestimated the impact of stomata because very few SLGCs have stomatal neighbours at this early developmental stage.

In this work we focused primarily on the influences coming from the immediate neighbourhood of an SLGC. It is also important to consider whether SLGC behaviour is influenced by factors operating at larger spatial scales. As a proxy for tissue-wide effects, we included each SLGC’s position in the leaf—the X- and Y-coordinates—in our original decision tree analysis (Fig. 1D; Supplemental Fig. 1A). Neither coordinate was predictive of SLGC behaviour, which indicates that the non-cell-autonomous factors that govern SLGC behaviour are primarily local.

We showed that the SPCH dynamics in dividing and differentiating cells bifurcate in early G1 (∼200 minutes after birth, Fig. 2), which raises the question of whether the decision to divide is made at this point. The dynamics of CDT1A, a replication licensing factor, appear to support this notion (Supplemental Fig. 2A). In dividing cells, CDT1A begins to accumulate in early G1, approximately 300 minutes after birth (Supplemental Fig. 2A). Since CDT1A accumulation is a hallmark of cell cycle progression (Desvoyes et al., 2019), these dynamics suggest that the decision to divide is made in early G1, and no later than ∼300 minutes after birth.

There is precedent for cell fate decisions in early G1. For example, human embryonic stem cells can only differentiate into endoderm if they receive TGF-β-Smad2/3 signals in early G1, when cyclin D levels are low (Pauklin and Vallier, 2013). Once cyclin D levels rise in late G1, the Smad2/3 proteins are phosphorylated, which prevents them from entering the nucleus and activating endoderm genes (Pauklin and Vallier, 2013). Likewise, the decision to undergo a formative or proliferative division in *Arabidopsis* roots is thought to occur in early G1 (Winter et al., 2024). Cells expressing low, transient levels of the transcription factors SCARECROW and SHORTROOT in early G1 are likely to divide formatively, rather than proliferatively (Winter et al., 2024).

Interestingly, in cells that might be the most closely related to SLGCs, the giant cells of the sepal epidermis, cell fate decisions are linked to a concentration threshold of the HD-ZIP transcription factor AtML1 in G2 (Meyer et al., 2017). Two elements of the Meyer study provide useful context and contrasts to our work. Giant cells are highly endoreplicated, and differentiation of SLGCs into pavement cells is also accompanied by endoreplication. Models of AtML1 function suggest that its G2 expression enables regulation of genes that promote endoreplication over mitotic cycles in giant cells (Meyer et al., 2017). Our finding that [SPCH] in early G1 is predictive of SLGC fate would suggest that endoreplication is a secondary consequence, rather than a cause of differentiation. Second, Meyer’s work emphasizes the cell-autonomous nature of the giant cell fate choice, which fits into their mechanical role in sepal shape. SLGCs, on the other hand, are sensitive to non-cell-autonomous factors, which enables them to carry out the stem-cell-like function of modulating leaf cell numbers and types in response to external inputs. Considered together, our quantitative imaging, statistical, simulation, and experimental approaches identified some of the key players in the SLGC decision, including cell size, SPCH activity, and cell-cell signaling. Our work shows how initial differences in SPCH levels are canalized by nuclear size and signaling to inform flexible cell fate decisions. It also highlights the existence of SPCH-independent pathways that link cell size and signaling to behaviour, which will be an important avenue of study moving forward.

## Methods

### Plant material and growth conditions

All *Arabidopsis* lines were in the Col-0 background (“wild-type”). Seeds were surface-sterilized by ethanol or chlorine gas (protocols based on Clough and Bent, 1998) and stratified for two days. Following stratification, they were grown vertically on half-strength Murashige and Skoog (MS) media with 0.8% or 1% agar for five days under long-day conditions (16 h light : 8 h dark at 22°C) and moderate intensity, full-spectrum light (110 µE).

Newly generated and previously reported lines are described in Supplemental Table 5. All transgenes used have been reported previously. Transgenic lines were generated by floral dip (Clough and Bent, 1998) and transgenic seedlings were selected on half-strength MS with the appropriate antibiotic (50 mg/L kanamycin or hygromycin).

### Image acquisition and image analysis

All confocal imaging experiments were performed on a Leica SP5 or Stellaris 8 confocal microscope with HyD detectors, a 40x NA1.1 water objective, image size of 1024 x 1024 pixels, and a digital zoom of 0.75-1x (unless otherwise specified). Only the abaxial surfaces of cotyledons were imaged. Raw Z-stacks were projected with Sum Slices in Fiji (Schindelin et al., 2012).

#### Wild-type time-lapses

To explore the relationship between birth size and SLGC behaviour, we captured time-lapses of 3-dpg, wild-type seedlings bearing a plasma membrane marker (*pATML1::mCherry-RCI2A* or *pATML1::YFP-RCI2A*) and a polarity marker (*pBRXL2::BRXL2-YFP*), a nuclear marker (*pATML1::H2B-mTFP*), or a cell cycle marker (PlaCCI; Desvoyes et al., 2020). Seedlings were mounted in a custom imaging chamber with half-strength MS solution (Davies and Bergmann, 2014) and imaged at 40 or 45-minute intervals for ∼16 hours. Cell size and shape were extracted from ilastik segmentations of the plasma membrane channel (Fig. 1C; Berg et al., 2019; Gong, Dale, Fung, Amador et al., 2023). Features “at birth” were measured from the first frame in which the newly formed cell plate was visible. After imaging, seedlings were returned to half-strength MS agar plates, where they were grown under long-day conditions (16 h light : 8 h dark at 22°C) and moderate intensity light (110 µE). Two days later, they were re-imaged to capture the developmental outcomes of SLGCs and their neighbours. Stringent quality controls to ensure that cells in imaged seedlings were not arrested meant that we only used 25% of all time-lapse experiments generated.

#### CDT1A intensities

To quantify CDT1A intensities, we enclosed each SLGC nucleus in the PlaCCI time-lapse described above (Desvoyes et al., 2020) in a circular ROI (area: 45.28 µm^2^) and measured the background-subtracted raw integrated density of CFP within each ROI.

#### SPCH reporter time-lapses

To quantify SPCH levels in SLGCs, we acquired time-lapses of 3-dpg seedlings expressing a plasma membrane marker (*pATML1::mCherry-RCI2A*) and a SPCH translational reporter (*pSPCH::SPCH-YFP* rescuing *spch-3*). Individuals were imaged as described above, except for one individual (#4), which was mounted on a slide with vacuum grease and imaged for 8 hours at 60-minute intervals. To quantify SPCH intensities, we segmented the plasma membrane channel using ilastik (Berg et al., 2019) and measured the background-subtracted raw integrated density of YFP within the cell boundaries of each SLGC.

Because our SPCH reporter line lacked a genetically encoded nuclear marker, we could not measure nuclear concentrations directly from our data. Instead, we stained the nuclei of 3-dpg seedlings with Hoechst (protocol described in Gong, Dale, Fung, Amador et al., 2023), segmented both the nuclear and genetically encoded plasma membrane signals using ilastik (Berg et al., 2019), and fitted a linear regression model to our ln-transformed cell and nuclear area measurements (α = 1.82 ± 0.14, β = 0.19 ± 0.032, *t* = 5.83, *P* = 3.42e-08; Supplemental Fig. 2B). Our model was not heteroscedastic (Breusch-Pagan test: χ^2^ = 0.034, *P* = 0.85), so we assumed that the distribution of residuals at any point along the fitted line could be modeled as a normal distribution *N*(6.45e-18, 0.027). For each cell in our SPCH dataset, we estimated nuclear area from this equation: *ln*(nuclear area) = 0.19 * *ln*(cell area) + 1.82 + ϵ, where ϵ ∼ *N*(6.45e-18, 0.027). We divided SPCH intensity by nuclear area to obtain nuclear concentrations.

#### Analysis of the contribution of bleaching to SPCH behaviours

To determine whether bleaching alone could account for the observed decline in SPCH intensities, we estimated bleaching rates per individual seedling. Using the Fiji plugin TrackMate (Ershov et al., 2022), we quantified the background-subtracted raw integrated density of YFP in each nucleus of each frame of the time-lapse. We regressed intensity on time (in hours) and divided the slope by the intercept to estimate a bleaching rate in % per hour. To compute the overall rates of decline in SPCH intensities, we fit an exponential decay function (*N*(t) = *N*_0_ * *e*^λt^) to the SPCH intensities in each cell from 0 to ∼200 minutes after birth and estimated the decay constant λ. We computed the overall rate of decline (% SPCH lost per hour) as 100% * (1 – e^λ^). Bleaching rates were low, ranging from 1.5-2.2% per hour, compared to mean rates of decline of 23-42% per hour (Supplemental Table 1).

#### SPCH++ time-lapses

To test whether large cells divide when supplied with enough SPCH, we captured time-lapses of *spch-3* seedlings expressing a plasma membrane marker (*pATML1::mCherry-RCI2A*) and a transgene *pSPCH::SPCH-YFP* that overproduces SPCH in its native domain. SPCH overproduction was verified through phenotypic analysis (an increase in asymmetric cell divisions; Fig. 3A) and fluorescence quantification (Supplemental Fig. 3D). We imaged three individuals: one individual was imaged in the custom time-lapse chamber (Davies and Bergmann, 2014) and the remaining two were imaged on slides with vacuum grease at 0, 3, and 6 hours, before being returned to half-strength MS plates (Gong, Dale, Fung, Amador et al., 2023). Two days later, they were re-imaged to capture cell behaviours. SLGCs grow very slowly (mean ± standard deviation: 1.46 ± 0.88% per hour), so this modified protocol only increased the error in birth size measurements due to cell growth from ∼1 to ∼3%, while enabling a larger number of individuals to be imaged simultaneously.

#### SPCH2-4A time course

Three *spch-3* seedlings expressing a plasma membrane marker (*pATML1::mCherry-RCI2A*) and *pSPCH::SPCH2-4A-YFP* (Davies and Bergmann, 2014) were imaged once at 3-dpg and again at 5-dpg, using 25X and 40X objectives. Cell sizes were extracted from ilastik segmentations of the plasma membrane channel at 3-dpg (Berg et al., 2019).

#### CYCD7-YFP time course

A seedling expressing a plasma membrane marker (*pATML1::mCherry-RCI2A*) and *pATML1::CYCD7-YFP* was imaged once at 3 dpg and again at 5 dpg. Cell sizes were extracted from ilastik segmentations of the plasma membrane channel at 3 dpg (Berg et al. 2019). In addition to imaging the entire cotyledon at a digital zoom of 0.75x (Supplemental Fig. 3A), we also imaged one region at 4x (Supplemental Fig. 3B).

#### Measuring fraction of the cell perimeter in contact with a signaling neighbour

We calculated the fraction of the cell perimeter in contact with a given signaling neighbour as

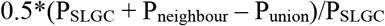

where Pslgc is the perimeter of the SLGC, Pneighbour is the perimeter of the signaling neighbour, and Punion is the perimeter of the union of the SLGC and the signaling neighbour. The total fraction of the cell perimeter in contact with a signaling neighbour is the sum of these fractions.

#### Statistical analysis

##### Classification and regression tree

To identify correlates of SLGC behaviour, we measured 15 features of SLGCs at birth (listed in Supplemental Fig. 1A) in a seedling expressing a plasma membrane marker (*pATML1::mCherry-RCI2A*) and a cell cycle marker (PlaCCI; Desvoyes et al., 2020). We fed our measurements into a classification and regression tree (CART), which we implemented in Python using the *scikit-learn* library (sklearn.tree module; Pedregosa et al., 2011). The CART method builds a decision tree by recursively partitioning cells along predictor axes into subsets that divide or differentiate. We first split our data into training and test sets (70:30) by randomly sampling without replacement. Next, we fit the CART algorithm to the training set, using Gini impurity as a measure of split quality, and applied cost complexity pruning to prevent overfitting. To build our final tree, we selected a cost complexity parameter value (α) that maximized testing accuracy.

We also specified a random forest classifier (sklearn.ensemble module), which controls overfitting by fitting 1,000 trees to various subsamples of the data and computing an average prediction accuracy. Feature importance scores were calculated as the normalized, total reduction of Gini impurity contributed by a given feature (Pedregosa et al., 2011).

##### Analyses in R

Mixed-effects models were specified using the *nlme* package (*v3.1-162*; Pinheiro and Bates, 2023) with predictors of interest as fixed effects and individual as a random effect. All other comparisons were made with unpaired Wilcoxon signed-rank tests, *Z-*tests for two proportions, or chi-squared tests for trend in proportions (*rstatix* package *v0.7.2*; Kassambara, 2024). Exponential, linear, and logistic models were fit with the *stats* package (*v4.3.1*; R Core Team, 2013), with individual as a blocking variable where appropriate.

#### Simulations

The lineage decision tree model was implemented in MATLAB 2021a and expands on the model reported in Gong, Dale, Fung, Amador et al. (2023). It is a stochastic, asynchronous rule-based model where SLGCs undergo birth, growth, SPCH degradation, and differentiation or division (Supplemental Fig. 5). All input parameters were derived from plant-specific, empirical distributions using MATLAB (see Supplemental Table 2 for fitted parameters). Cell sizes were rounded to the nearest integer µm^2^. Nuclear sizes were estimated as described above: *ln*(nuclear area) = 0.19 * *ln*(cell area) + 1.82 + ϵ, where ϵ ∼ *N*(6.45e-18, 0.027).

The starting sizes, SPCH intensities, and numbers of signaling neighbours of 1,000 mother cells were randomly drawn from gamma, normal, and Poisson distributions, respectively. The cells then divided with an asymmetry drawn from a beta distribution with a noise factor ±0.05 drawn from a uniform distribution, each forming a smaller meristemoid and a larger SLGC. Based on Supplemental Fig. 6, each SLGC was assumed to inherit two-thirds of its mother’s SPCH intensity and to have one signaling neighbour more than its mother (i.e. the newly generated sister meristemoid). SPCH degradation rates (decay constants, λ) were randomly drawn from four possible exponential distributions fit to the following:

- Neighbour-based degradation: degradation rates were split by the number of signaling neighbours (1 vs. 2+ neighbours)
- Size-based degradation: degradation rates were calculated on a per-micron basis
- Neighbour- and size-based degradation: degradation rates were calculated on a per-micron basis and split by the number of signaling neighbours
- Random (neighbour- and size-independent): degradation rates were pooled

The probability of division was determined based on a cell’s size, [SPCH] at the dip, number of signaling neighbours, and/or their interaction terms, using multiple logistic parameters estimated via the *stats* package in R (*v4.3.1*; R Core Team, 2013; Supplemental Table 3).

Model selection was performed by simulating across modes of SPCH degradation (random, neighbour-based, size-based, or neighbour- and size-based) and modes of fate determination (including linear and interaction terms of cell size, [SPCH], and the number of signaling neighbours) in a factorial manner. Simulations were run for one generation. Model selection occurred in three steps. First, the sizes and [SPCH] of dividing and differentiating cells were compared to those of the data using “two one-sided tests” (TOST) equivalence testing. The null hypothesis in TOST equivalence testing is that there is a difference in populations greater than the effect size of interest. Due to our experimental sample size, we chose an effect size of one standard deviation (Lakens, 2017). We used Welch’s *t*-tests for unequal sample sizes with the Satterthwaite correction. A significance threshold of 0.05 was used to determine equivalence.

Second, two-sample *t*-tests were run to check if there were significant differences in the sizes and [SPCH] of cells that divided vs. differentiated. A threshold of 0.05 was used to determine significance. Finally, the sum of squared errors (SSE) was calculated to assess the fit of each model to the proportion of cells that divided given the number of signaling neighbors. The Akaike information criterion (AIC) was calculated for the total SSE across individuals using the following formula: 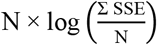, where *N* is the total number of cells in the data (262). Models with additional parameters in the fate-determining logistic were penalized by 2 AIC points for each additional factor or interaction term.

## Notes

### Competing Interest Statement

The authors have declared no competing interest.

