## Supplemental Figures 1-7; Tables 1-6 for "Multi-scale dynamics influence the division potential of stomatal lineage ground cells in *Arabidopsis*"

### Contents:

**Supplemental Figures S1-S7**

**Supplemental Tables S1-S6**

A

Features tested for predictive value in decision tree analysis

- Time of birth
- X, Y-coordinates
- Birth area of SLGC ($\mu m^2$)
- Birth area of sister meristemoid ($\mu m^2$)
- Asymmetry of birth division =
  (SLGC – meristemoid area)/(SLGC + meristemoid area)
- Major axis: length of major axis
- Minor axis: length of minor axis
- Circularity = $4\pi$*area/perimeter$^2$
- Aspect ratio = major axis/minor axis
- Roundness = 4*area/($\pi$*(major axis)$^2$)
- Solidity = area/convex area
- Mother identity (meristemoid or SLGC)
- Sister behaviour
  - divided asymmetrically (ACD)
  - differentiated
  - neither divided nor differentiated
- Number of adjacent, non-sister stomatal precursors
  - meristemoids, GMCs, or stomata

B

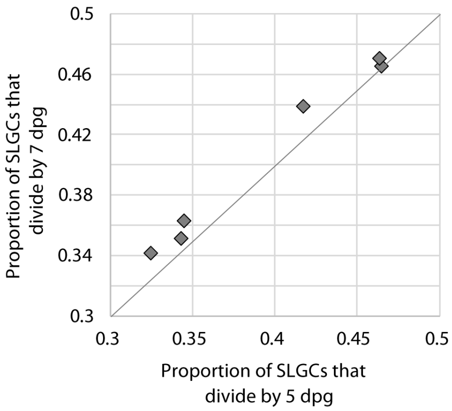


C

| Effective α | Nodes | Maximum depth | Accuracy |
|---|---|---|---|
| 0 | 13 | 4 | 73.7% |
| 0.03488372 | 9 | 3 | 68.4% |
| 0.06659619 | 7 | 3 | 68.4% |
| 0.07475083 | 5 | 2 | 78.9% |
| 0.07539716 | 3 | 1 | 73.7% |
| 0.09423472 | 1 | 0 | 73.7% |

D

| Feature | Importance score |
|---|---|
| Birth size | 0.122385 |
| Time of birth | 0.121212 |
| Major axis | 0.116688 |
| Circularity | 0.090541 |
| Minor axis | 0.077721 |
| Y-coordinate | 0.071104 |
| Solidity | 0.066173 |
| Sister meristemoid size | 0.063194 |
| Division asymmetry | 0.060259 |
| X-coordinate | 0.056747 |
| Roundness | 0.053900 |
| Aspect ratio | 0.050392 |
| Sister behaviour<br>• ACD<br>• differentiation<br>• neither ACD nor differentiation | <br>• 0.020444<br>• 0.003801<br>• 0.014964 |
| Number of adjacent, non-sister stomatal precursors | 0.007750 |
| Mother identity | 0.002725 |

**Supplemental Figure 1. Additional details on the time-lapse and decision tree analyses.** (A) List of features at birth that were tested for predictive value in the decision tree analysis. (B) Proportion of SLGCs born on 3-dpg that divided by 7-dpg vs. proportion of cells that divided by 5-dpg. The vast majority of cells divided between 3- and 5-dpg. (C) The number of nodes, maximum depth, and accuracy for different cost complexity parameter values (effective α). The parameter value that maximized testing accuracy is highlighted in yellow. (D) The feature importance scores (total reduction in Gini impurity) for all features tested. These values were derived from a random forest classifier with 1,000 trees.

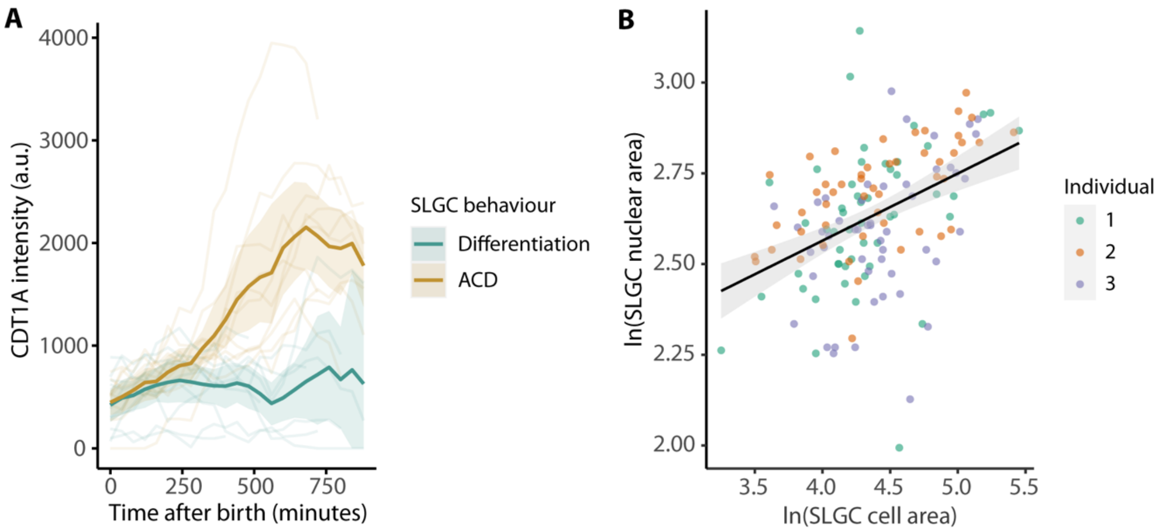


**Supplemental Figure 2. CDT1A dynamics after birth and the relationship between cell and nuclear areas.** (A) CDT1A-CFP (from the PlaCCI reporter line; Desvoyes et al., 2020) intensities over time, coloured by behaviour. In dividing cells (ACD), CDT1A-CFP increased ~300 minutes after birth. Lines and bands are means and bootstrapped 95% confidence intervals. *N* = 20 cells, 10 cells per behaviour. (B) SLGC nuclear area vs. cell area. Axes are ln-transformed. The black line and grey band are linear model predictions and 95% confidence intervals. *N* = 3 individuals, 50 cells per individual.

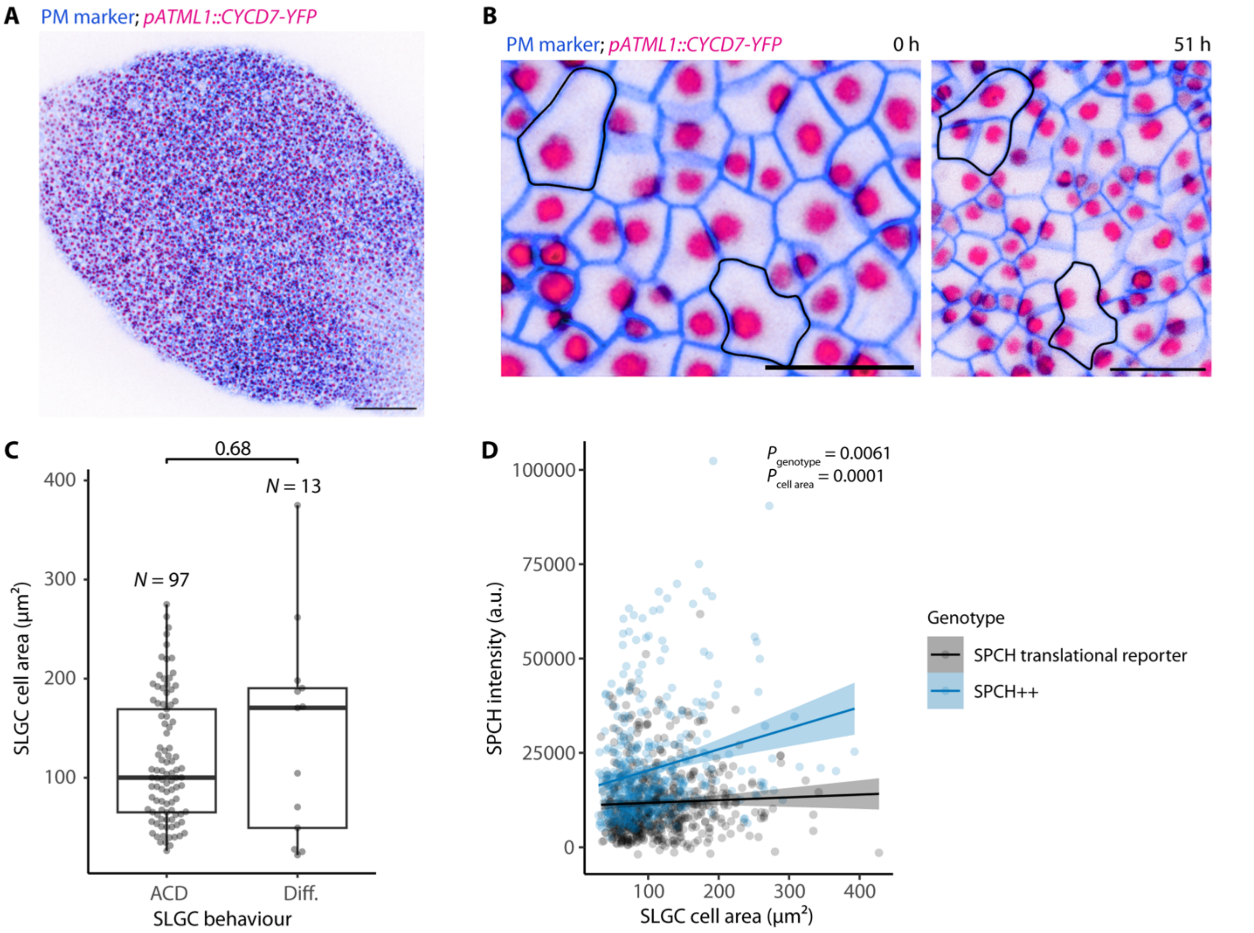


**Supplemental Figure 3. The epidermis-specific expression of a D-type cyclin induces large SLGCs to divide.** (A) Micrograph of a 3-dpg cotyledon expressing a plasma membrane (PM) marker and the D-type cyclin CYCD7;1 (CYCD7) under the epidermis-specific *ATML1* promoter (*pATML1::CYCD7;1-YFP*). Scale bar: 100 µm. (B) The *pATML1::CYCD7;1-YFP* construct was present in all epidermal cells at 3 dpg (left). The same region 51 hours later (right). Large, dividing SLGCs are outlined in black. Scale bar: 20 µm. (C) Cell areas of SLGCs that divided (ACD) or differentiated (Diff.) in a 3-dpg cotyledon expressing *ATML1p::CYCD7;1-YFP*. The *P*-value is calculated by a Wilcoxon rank-sum test. $N = 110$ cells. (D) SPCH intensity vs. cell area in *spch-3* cotyledons expressing the SPCH translational reporter or a *pSPCH::SPCH-YFP* transgene that overproduces SPCH (SPCH++). *P*-values are from a mixed-effects model with cell area and genotype as fixed effects and individual as a random effect. $N$ = 4-6 individuals per genotype, 85-154 cells per individual.

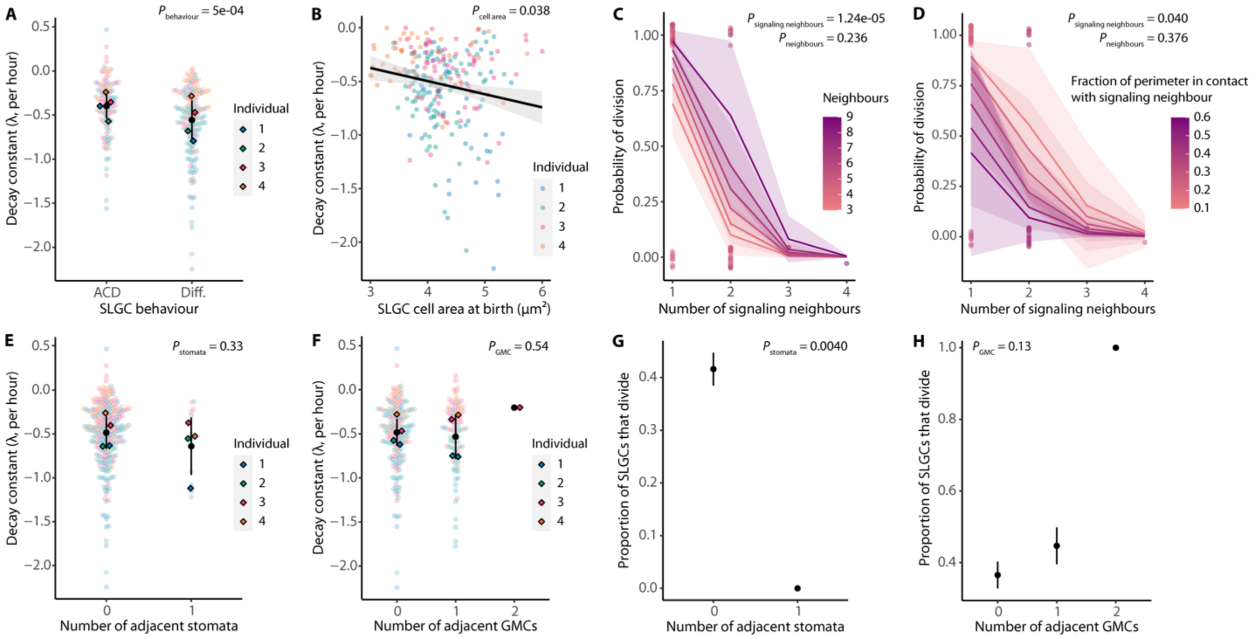


**Supplemental Figure 4. The effect of signaling neighbours on SPCH degradation rate is not explained by the number of adjacent stomata or GMCs.** (A) The decay constant (how fast SPCH is degraded) in dividing (ACD) or differentiating (Diff.) cells. The more negative the constant, the higher the degradation rate. (B) The decay constant vs. cell area at birth (ln-transformed) among cells where SPCH levels declined. The black line and grey band are linear model predictions and 95% confidence intervals. (C-D) Multiple logistic regression of the probability of division on the number of signaling neighbours and the total number of neighbours (C) or the fraction of the cell perimeter in contact with a signaling neighbour (D). Lines and bands are logistic model predictions and standard errors. *P*-values are from the logistic regressions. $N = 75$ cells. (E-F) The decay constant by the number of adjacent stomata (E) or GMCs (F). (G-H) The proportion of SLGCs that divided by the number of adjacent stomata (G) or GMCs (H). (A,E-H) Black circles and lines are means and standard deviations. (A,B,E-H) *P*-values are from mixed-effects models with individual as a random effect (A-B,E-F) or chi-squared tests for trend in proportions (G-H). $N = 4$ individuals, 50-75 cells per individual.

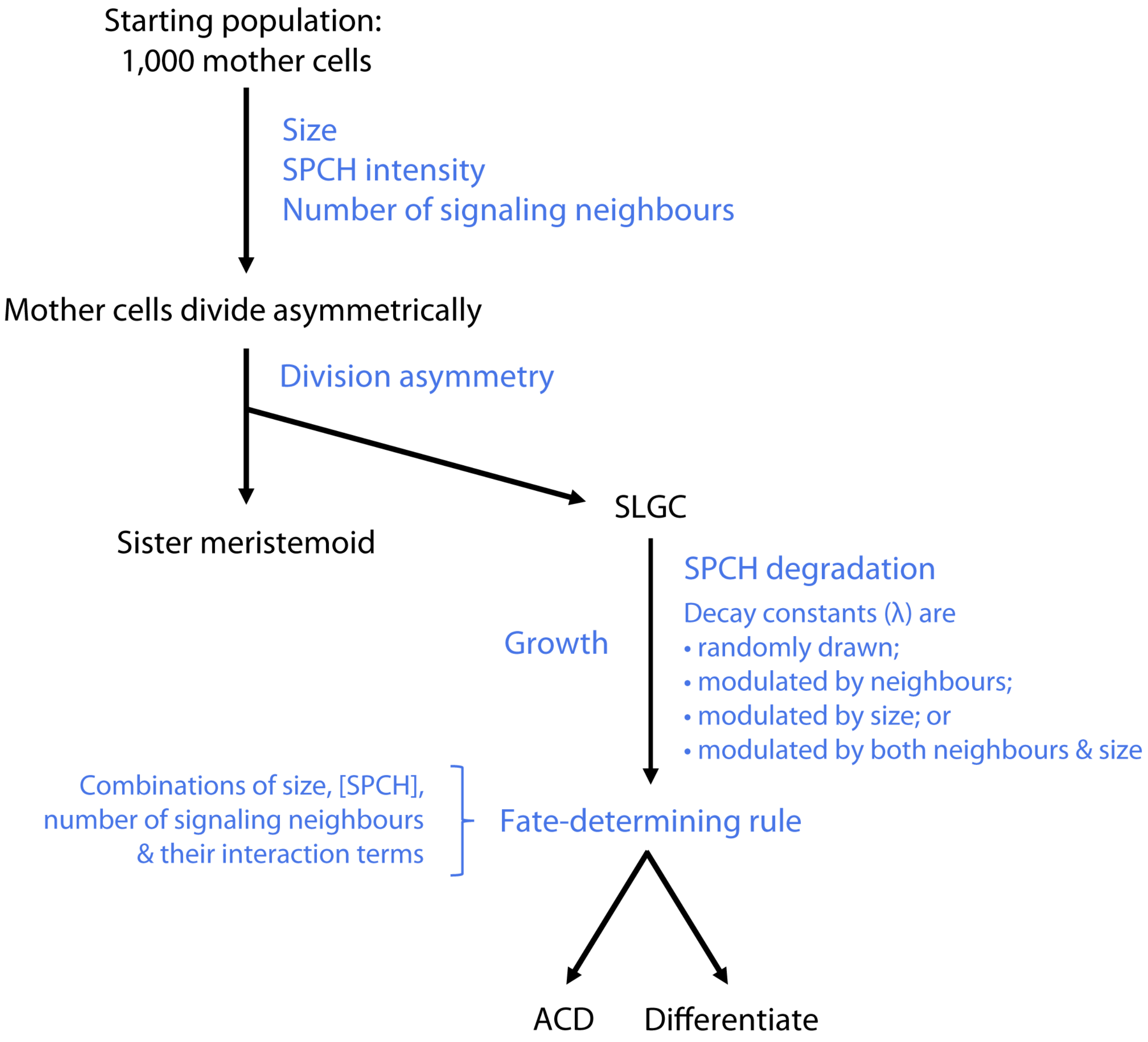


**Supplemental Figure 5. Diagram of the lineage decision tree model.** The model begins with 1,000 mother cells, each with a randomly drawn starting size, SPCH intensity, and number of signaling neighbours (derived from empirical data; see Supplemental Tables 2 and 3 for fitted parameters). The mother cells divide with a randomly drawn asymmetry, forming a smaller meristemoid and a larger SLGC. Based on experimental measurements (Supplemental Figure 6), each SLGC is assumed to inherit two-thirds of its mother's SPCH intensity and to have one signaling neighbour more than its mother. To identify the model that best recapitulated the data, we simulated across modes of SPCH degradation and modes of fate determination in a factorial manner.

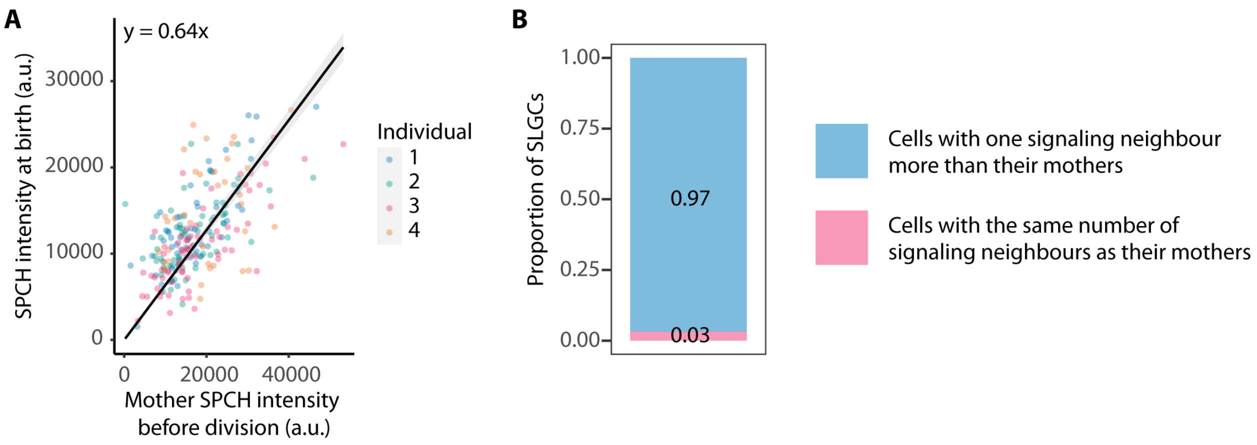


**Supplemental Figure 6. SLGCs inherit approximately two-thirds of their mothers' SPCH intensities and have one signaling neighbour more than their mothers.** (A) SPCH intensity at birth vs. mother SPCH intensity before division. The black line and grey band are linear model predictions and 95% confidence intervals. *N* = 4 individuals, 50-75 cells per individual. (B) The proportion of SLGCs with one signaling neighbour more than their mothers (blue) and the same number of signaling neighbours as their mothers (pink). *N* = 2 individuals, 50-75 cells per individual.

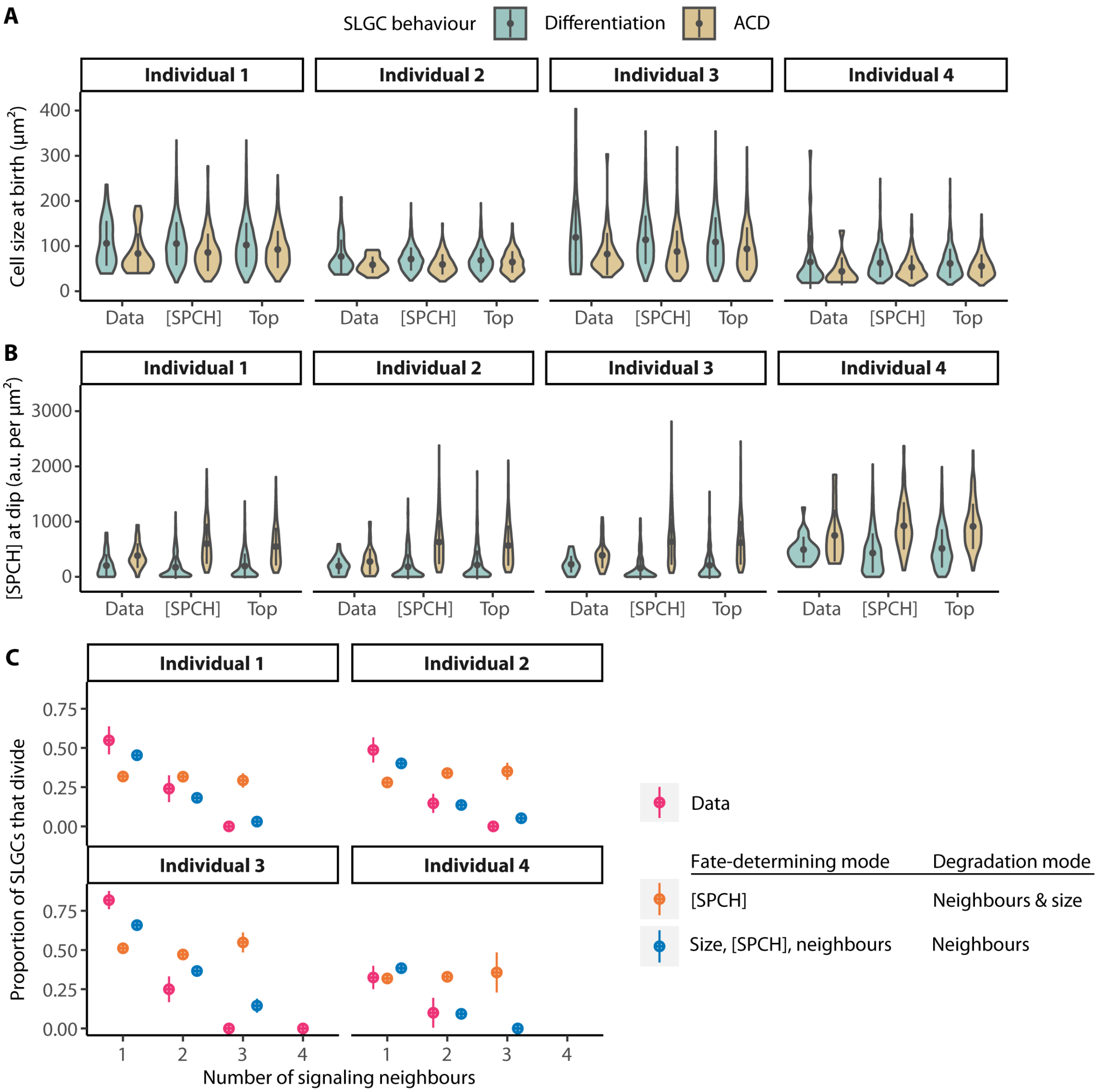


**Supplemental Figure 7. A fate-determining program with size, [SPCH], and signaling neighbours outperforms one with [SPCH] alone.** (A-B) The birth sizes (A) and [SPCH] at the dip (B) of dividing (ACD) and differentiating SLGCs in the data, compared to those in the highest-ranking [SPCH] model ("[SPCH]") and the top model ("Top"). In the highest-ranking [SPCH] model, degradation rates scale with both signaling neighbours and size, and only [SPCH] is considered in the fate-determining process. In the top model, degradation rates scale with signaling neighbours alone, and size, [SPCH], and signaling neighbours are all considered in the fate-determining process. (C) The proportion of SLGCs that divided given the number of signaling neighbours in the data (pink), highest-ranking [SPCH] model (orange), and top model (blue). (A-C) Circles and lines are means and standard deviations. Data: $N = 4$ individual plants, 50-75 cells per individual. Models: $N = 4$ individual plants, 1,000 cells per individual.

**Supplemental Table 1. Estimates of SPCH-YFP bleaching rates, compared to overall rates of decline in SPCH-YFP intensities from 0 to ~200 minutes after birth.**

| Individual | Slope | Intercept | Bleaching rate (slope/intercept, per hr) | Mean decay constant (λ) | Overall rate of decline (% SPCH lost per hr) |
|---|---|---|---|---|---|
| 1 | -235.61 | 14125 | -1.67% | -0.649 | -42.4% |
| 2 | -257.20 | 13163 | -1.95% | -0.640 | -44.0% |
| 3 | -339.80 | 15290 | -2.22% | -0.404 | -30.2% |
| 4 | -212.90 | 14647 | -1.45% | -0.280 | -22.7% |

**Supplemental Table 2. Fitted parameters for simulations.**
*Parameters for neighbour-based degradation. **Parameters for neighbour- and size-based degradation.

| Variable | Distribution | Fitted parameters | | | |
|---|---|---|---|---|---|
| | | **Individual 1** | **Individual 2** | **Individual 3** | **Individual 4** |
| Division asymmetry | Beta | α = 6.796<br>β = 14.278 | α = 9.706<br>β = 18.582 | α = 5.696<br>β = 13.385 | α = 5.850<br>β = 10.253 |
| Number of signaling neighbours of mother cell | Poisson | λ = 0.597 | λ = 0.507 | λ = 0.467 | λ = 0.200 |
| Mother cell size before division (μm$^2$) | Gamma | α = 5.033<br>β = 28.878 | α = 8.168<br>β = 12.561 | α = 3.668<br>β = 38.219 | α = 2.822<br>β = 28.414 |
| Mother SPCH intensity before division (a.u.) | Normal | μ = 2.688<br>σ = 1.025 | μ = 2.398<br>σ = 0.776 | μ = 2.267<br>σ = 1.121 | μ = 2.749<br>σ = 1.076 |
| SLGC growth rate (%/hr) | Mean | 1.013 | 1.011 | 1.010 | 1.006 |
| Randomly drawn degradation rates (λ) | Exponential | μ = 0.6644 | μ = 0.6410 | μ = 0.4193 | μ = 0.2823 |
| Degradation rates (λ) for cells with 1 neighbour* | Exponential | μ = 0.5417 | μ = 0.5878 | μ = 0.3429 | μ = 0.2494 |
| Degradation rates (λ) for cells with 2+ neighbours* | Exponential | μ = 0.7755 | μ = 0.7009 | μ = 0.5098 | μ = 0.4005 |
| Size-based, per-micron degradation rates | Exponential | μ = 0.0079 | μ = 0.102 | μ = 0.0056 | μ = 0.0068 |
| Size-based, per-micron degradation rates for cells with 1 neighbour** | Exponential | μ = 0.0075 | μ = 0.108 | μ = 0.0049 | μ = 0.0072 |
| Size-based, per-micron degradation rates for cells with 2+ neighbours** | Exponential | μ = 0.0082 | μ = 0.0096 | μ = 0.0063 | μ = 0.0052 |
| Duration of degradation (hr) | Mean | 2.5574 | 2.7295 | 2.6667 | 2.2917 |

**Supplemental Table 3. Multiple logistic regression parameters for modes of fate determination.**

| Model | Terms | Estimate (β) |
|---|---|---|
| Size | Intercept | 0.551870 |
| | Size at dip | -0.011532 |
| | Individual 2 | -0.485926 |
| | Individual 3 | 0.891775 |
| | Individual 4 | -0.819261 |
| [SPCH] | Intercept | -1.5175689 |
| | [SPCH] at dip | 0.0033391 |
| | Individual 2 | -0.0210154 |
| | Individual 3 | 0.7971070 |
| | Individual 4 | -1.3871696 |
| Neighbours | Intercept | 2.3762 |
| | Signaling neighbours | -2.0042 |
| | Individual 2 | -0.3592 |
| | Individual 3 | 0.8574 |
| | Individual 4 | -1.0107 |
| Size + [SPCH] | Intercept | -0.4648068 |
| | Size at dip | -0.0113864 |
| | [SPCH] at dip | 0.0034161 |
| | Individual 2 | -0.2722333 |
| | Individual 3 | 0.8715108 |
| | Individual 4 | -1.8741598 |
| [SPCH] + neighbours | Intercept | 1.3555498 |
| | [SPCH] at dip | 0.0030930 |
| | Signaling neighbours | -1.9100477 |
| | Individual 2 | -0.1847612 |
| | Individual 3 | 0.8306865 |
| | Individual 4 | -2.0130550 |
| Size + neighbours | Intercept | 2.745461 |
| | Size at dip | -0.005924 |
| | Signaling neighbours | -1.855097 |
| | Individual 2 | -0.528401 |
| | Individual 3 | 0.857530 |
| | Individual 4 | -1.233415 |
| [SPCH] + neighbours + [SPCH]*neighbours | Intercept | 1.4656494 |
| | [SPCH] at dip | 0.0027066 |
| | Signaling neighbours | -1.9958963 |
| | [SPCH] at dip*signaling neighbours | 0.0002894 |
| | Individual 2 | -0.1856469 |
| | Individual 3 | 0.8382843 |
| | Individual 4 | -1.9789828 |
| Size + neighbours + size*neighbours | Intercept | 1.876672 |
| | Size at dip | 0.003938 |
| | Signaling neighbours | -1.209844 |
| | Size at dip*signaling neighbours | -0.007204 |
| | Individual 2 | -0.496369 |
| | Individual 3 | 0.878791 |
| | Individual 4 | -1.145140 |
| Size + [SPCH] + neighbours | Intercept | 1.7030774 |
| | Size at dip | -0.0058269 |
| | [SPCH] at dip | 0.0030755 |
| | Signaling neighbours | -1.7579984 |
| | Individual 2 | -0.3388473 |
| | Individual 3 | 0.8447111 |

| | | |
|---|---|---|
| | Individual 4 | -2.2166096 |
| Size + [SPCH] + neighbours + size*neighbours + [SPCH]*neighbours | Intercept<br>Size at dip<br>[SPCH] at dip<br>Signaling neighbours<br>Size at dip*signaling neighbours<br>[SPCH] at dip*signaling neighbours<br>Individual 2<br>Individual 3<br>Individual 4 | 1.0352947<br>0.0034542<br>0.0025846<br>-1.2775606<br>-0.0066062<br>0.0003713<br>-0.3124349<br>0.8679143<br>-2.0895315 |
| Size + neighbours + size*[SPCH] | Intercept<br>Size at dip<br>Signaling neighbours<br>Size at dip*[SPCH] at dip<br>Individual 2<br>Individual 3<br>Individual 4 | 2.713<br>-0.01677<br>-1.736<br>0.00003119<br>-0.3936<br>0.8582<br>-1.770 |

**Supplemental Table 4. Goodness-of-fit of each model to the proportion of SLGCs that divided given the number of signaling neighbours.**

The top model, with the lowest AIC score, is highlighted in orange. Only models that passed equivalence and significance tests (see Methods) were evaluated.

| **Model** | | **Sum of squared errors (SSE), by individual** | | | | **Akaike information criterion (AIC)** |
|---|---|---|---|---|---|---|
| **Mode of fate determination** | **Mode of SPCH degradation** | **1** | **2** | **3** | **4** | |
| Size | Random | Failed equivalence and/or significance tests (see Methods) | | | | N/A |
| | Neighbour | | | | | |
| | Size | | | | | |
| | Neighbour & size | | | | | |
| [SPCH] | Random | Failed equivalence and/or significance tests | | | | N/A |
| | Neighbour | | | | | |
| | Size | 0.153 | 0.176 | 0.465 | 0.106 | -645.599 |
| | Neighbour & size | 0.150 | 0.182 | 0.444 | 0.122 | -645.860 |
| Neighbours | Random | Failed equivalence and/or significance tests | | | | N/A |
| | Neighbour | | | | | |
| | Size | | | | | |
| | Neighbour & size | | | | | |
| Size + [SPCH] | Random | 0.155 | 0.188 | 0.305 | 0.186 | -652.308 |
| | Neighbour | 0.109 | 0.147 | 0.423 | 0.159 | -651.709 |
| | Size | 0.133 | 0.186 | 0.488 | 0.158 | -635.715 |
| | Neighbour & size | 0.129 | 0.197 | 0.457 | 0.189 | -634.769 |
| [SPCH] + neighbours | Random | Failed equivalence and/or significance tests | | | | N/A |
| | Neighbour | | | | | |
| | Size | 0.020 | 0.015 | 0.103 | 0.001 | -856.738 |
| | Neighbour & size | 0.015 | 0.016 | 0.077 | 0.005 | -878.807 |
| Size + neighbours | Random | Failed equivalence and/or significance tests | | | | N/A |
| | Neighbour | | | | | |
| | Size | | | | | |
| | Neighbour & size | | | | | |
| [SPCH] + neighbours + [SPCH]*neighbours | Random | Failed equivalence and/or significance tests | | | | N/A |
| | Neighbour | | | | | |
| | Size | 0.021 | 0.017 | 0.119 | 0.001 | -839.625 |
| | Neighbour & size | 0.017 | 0.019 | 0.088 | 0.021 | -849.594 |
| Size + neighbours + size*neighbours | Random | Failed equivalence and/or significance tests | | | | N/A |
| | Neighbour | | | | | N/A |
| | Size | | | | | N/A |
| | Neighbour & size | | | | | N/A |
| Size + [SPCH] + neighbours | Random | Failed equivalence and/or significance tests | | | | N/A |
| | Neighbour | 0.013 | 0.010 | 0.060 | 0.004 | -907.359 |
| | Size | 0.024 | 0.017 | 0.121 | 0.000 | -836.755 |
| | Neighbour & size | 0.020 | 0.019 | 0.090 | 0.005 | -858.573 |
| Size + [SPCH] + neighbours + size*neighbours + [SPCH]*neighbours | Random | Failed equivalence and/or significance tests | | | | N/A |
| | Neighbour | | | | | |
| | Size | | | | | |
| | Neighbour & size | | | | | |
| Size + neighbours + size*[SPCH] | Random | Failed equivalence and/or significance tests | | | | N/A |
| | Neighbour | | | | | |
| | Size | | | | | |
| | Neighbour & size | 0.023 | 0.021 | 0.080 | 0.000 | -866.731 |

**Supplemental Table 5. Newly generated and previously reported lines.**

| Genetic line | Sources |
|---|---|
| PlaCCI cell cycle reporter<br>*pATML1::mCherry-RCI2A* | This manuscript. We introduced a plasma membrane marker into the cell cycle marker line generated by Desvoyes et al. (2020). |
| *pBRXL2::BRXL2-YFP*<br>*pATML1::mCherry-RCI2A* | Gong et al. (2021a) |
| *pATML1::H2B-mTFP*<br>*pATML1::mCitrine-RCI2A* | Robinson et al. (2018) |
| *spch-3*<br>*pSPCH::SPCH-YFP*<br>*pATML1::mCherry-RCI2A* | Lopez-Anido et al. (2021) |
| *spch-3*<br>*pSPCH::gSPCH-YFP* (SPCH++)<br>*pATML1::mCherry-RCI2A* | This manuscript. We introduced a plasma membrane marker into a genomic SPCH reporter line that overproduces SPCH (Vatén et al., 2018). |
| *pATML1::CYCD7;1-YFP*<br>*pATML1::mCherry-RCI2A* | This manuscript. We transformed a wild-type line bearing a plasma membrane marker with the *pATML1::CYCD7;1-YFP* construct generated by Weimer et al. (2018). |
| *spch-3*<br>*pSPCH::SPCH2-4A-YFP*<br>*pATML1::mCherry-RCI2A* | This manuscript. We introduced a plasma membrane marker into the *pSPCH::SPCH2-4A-YFP; spch-3* generated by Davies & Bergmann (2014). |

**Supplemental Table 6. Detailed statistics reported in each figure.**

| **Figure panel (listed in order described in main text)** | **Additional statistical details** |
|---|---|
| Fig. 1E | $\beta = -35.68 \pm 8.01\ \mu m^2$, df = 199, $t = -4.45$, $P = 1e\text{-}04$ |
| Fig. 2C | $\beta = 2082.43 \pm 634.28$ a.u., df = 256, $t = 3.28$, $P = 0.0012$ |
| Fig. 2F | $\beta = 0.039 \pm 0.049$, df = 256, $t = 0.79$, $P = 0.43$ |
| Fig. 2G | $\beta = -0.16 \pm 0.053$, df = 256, $t = -2.99$, $P = 0.0031$ |
| Fig. 3A | $Z = 5.003$, $P < 0.00001$ |
| Fig. 3B | $\beta = 5.09 \pm 13.72\ \mu m^2$, df = 4, $t = 0.37$, $P = 0.73$ |
| Fig. S4A | $\beta = 0.16 \pm 0.046$, df = 252, $t = 3.52$, $P = 5e\text{-}04$ |
| Fig. S4B | $\beta = -0.089 \pm 0.043$, df = 243, $t = -2.09$, $P = 0.038$ |
| Fig. 4B | $\beta = 37.04 \pm 5.11\ \mu m^2$, df = 256, $t = 7.25$, $P < 2.2e\text{-}16$ |
| Fig. 4C | $\chi^2 = 37.62$, df = 1, $P = 8.59e\text{-}10$ |
| Fig. 4D-E′ | $\beta = -0.13 \pm 0.038\ h^{-1}$, df = 252, $t = -3.49$, P = 0.0006 |
| Fig. 4F-G′ | $\beta = -0.071 \pm 0.033\ h^{-1}$, df = 252, $t = -2.17$, P = 0.031 |
| Fig. 4H | $\beta = 33.61 \pm 4.18\ \mu m^2$, df = 257, $t = 8.05$, $P < 2.2e\text{-}16$ |
| Fig. 4I | $\chi^2 = 27.73$, df = 1, $P = 1.39e\text{-}07$ |
| Fig. S4G | $Z = 2.88$, $P = 0.0040$ |
| Fig. S4H | $\chi^2 = 2.31$, df = 1, $P = 0.129$ |
| Fig. 5B | $\beta = 21.42 \pm 8.87\ \mu m^2$, df = 236, $t = 2.41$, $P = 0.017$ |
| Fig. 6B | $\beta = 157.62 \pm 29.37$, df = 245, $t = 5.37$, $P < 2.2e\text{-}16$ |
| Fig. 6C | $\beta_{\text{nuclear size}} = -1.16e\text{-}01 \pm 5.48e\text{-}02$, $Z = -2.11$, $P = 0.035$ |
